# The perinuclear ER scales nuclear size independently of cell size in early embryos

**DOI:** 10.1101/818724

**Authors:** Richik Nilay Mukherjee, Jérémy Sallé, Serge Dmitrieff, Katherine Nelson, John Oakey, Nicolas Minc, Daniel L. Levy

## Abstract

**SUMMARY:** Nuclear size plays pivotal roles in gene expression, embryo development, and disease. A central hypothesis in organisms ranging from yeast to vertebrates is that nuclear size scales to cell size. This implies that nuclei may reach steady state sizes set by limiting cytoplasmic pools of size-regulating components. By monitoring nuclear dynamics in early sea urchin embryos, we found that nuclei undergo substantial growth in each interphase, reaching a maximal size prior to mitosis that declined steadily over the course of development. Manipulations of cytoplasmic volume through multiple chemical and physical means ruled out cell size as a major determinant of nuclear size and growth. Rather, our data suggest that the perinuclear endoplasmic reticulum, accumulated through dynein activity, serves as a limiting membrane pool that sets nuclear surface growth rate. Partitioning of this local pool at each cell division modulates nuclear growth kinetics and dictates size scaling throughout early development.

## INTRODUCTION

Cell sizes vary dramatically throughout biology, in different cell types and organisms as well as during early development when reductive cell divisions occur without growth. A fundamental question in cell biology is how organelle size is adapted to cell size, a phenomenon referred to as organelle size scaling (Levy and Heald, 2012; Chan and Marshall, 2010). One central hypothesis is that organelle size scaling results from limiting cytoplasmic pools of diffusible growth-regulating material. As cells become smaller, this pool is more rapidly depleted, slowing organelle growth and reducing final organelle size (Goehring and Hyman, 2012). Studies of size scaling of mitotic spindles and centrosomes provide support for this general class of model (Lacroix, et al., 2018; Decker, et al., 2011). Alternative models include time-dependent regulation of organelle growth matched directly or indirectly with the timing of cell growth and division (Goehring and Hyman, 2012). On conceptual grounds, these two classes of models resemble important debates between timer and sizer models regulating cell size at division (Cadart, et al., 2018; Facchetti, et al., 2017).

The nucleus is one organelle that exhibits exquisite size scaling both during development and between species. Pioneering studies from Conklin and Wilson in the early 1900’s first reported on the scaling of cell and nuclear size (Wilson, 1925; Conklin, 1912), a relationship which has now been largely validated in numerous contexts ranging from single yeast cells to multicellular embryos and tissues (Vukovic, et al., 2016a; Levy and Heald, 2012). Scaling of nuclear size to cell size impacts the nuclear-to-cytoplasmic (N/C) volume ratio, with critical functional implications in development and disease. For instance, altering the N/C ratio in *Xenopus* embryos affects developmental timing and zygotic genome activation (Jevtic and Levy, 2015; Newport and Kirschner, 1982). Cancer cells with enlarged nuclei almost always represent more aggressive metastatic disease, and graded increases in nuclear size have long been used by pathologists to diagnose and stage almost all types of cancer, often independently of gross changes in ploidy (Jevtic and Levy, 2014). As such, addressing mechanisms that contribute to nuclear size scaling and links to developmental programs and tissue context remain outstanding problems in biology and medicine.

Mechanisms that regulate nuclear morphology most commonly invoke a role for structural elements that shape the nuclear envelope (NE) (Vukovic, et al., 2016a). The NE is composed of two lipid bilayers. The outer nuclear membrane is continuous with the endoplasmic reticulum (ER), and several studies have implicated membrane synthesis and NE-ER connections in the assembly and morphology of the NE (Golden, et al., 2009; Anderson and Hetzer, 2008; Anderson and Hetzer, 2007). The inner nuclear membrane is lined by the nuclear lamina, composed of lamin intermediate filaments and lamin-associated proteins. In addition to providing mechanical support to the NE and regulating chromatin organization, the lamina also influences nuclear size (Jevtic, et al., 2015; Levy and Heald, 2010). Nuclear pore complexes (NPCs) inserted into the NE are conduits for nucleocytoplasmic transport. Proteins containing a nuclear localization signal (NLS) are targeted to the NPC for nuclear import through association with importins. Nuclear shape and size depend on nuclear import of cargos such as the lamins (Newport, et al., 1990), and the levels and localization of several nuclear import factors have been shown to regulate nuclear size (Brownlee and Heald, 2019; Kume, et al., 2017; Ladouceur, et al., 2015; Levy and Heald, 2010). Thus, in principle, nuclear size could emerge as a result of a variety of activities, but how these activities might titrate nuclear size in development and disease remains poorly understood.

The hypothesis that nuclear size is determined by cell size implies that nuclei can rapidly reach a steady state size. However, the constant scaling of nuclear size to cell size in growing cells such as yeasts, which grow larger throughout interphase, relies on steady growth of nuclei (Jorgensen, et al., 2007; Neumann and Nurse, 2007). Whether cell and nuclear growth are simply concomitant, occurring at a similar expansion rate, or strongly coupled at each time point through size sensing mechanisms remains unclear (Goehring and Hyman, 2012). Given the complications introduced by cell growth, blastomere cells of early cleaving embryos, which are marked by dramatic cell size reductions with no growth, serve as a powerful context to test mechanisms of nuclear size scaling. However, in such systems, nuclear scaling has been mostly addressed in fixed embryos or using *in vitro* extracts, where it is not possible to examine how dynamic nuclear assembly, nuclear growth, and cell division influence nuclear size scaling. In addition, methods lack to systematically alter blastomere size to discern if nuclear size reductions in early development simply occur concomitantly with, or are actively regulated by, changes in cell size. Thus, the basic conceptual elements that regulate nuclear size and their connection to cell size and developmental programs remain to be established *in vivo*.

By exploiting the transparency and facile manipulation of developing sea urchin embryos, we here revisit mechanisms of nuclear size scaling. We find that nuclei of early blastomeres exhibit significant surface growth and that their maximum size is limited by the time to nuclear envelope breakdown (NEB) prior to mitosis. Steady state nuclear sizes are only reached at the 16-cell stage and beyond. By blocking cell cycle progression or altering cytoplasmic volumes, we demonstrate uncoupling of cell and nuclear sizes. Rather, our data support a model in which the perinuclear ER (pER) acts as a local limiting membrane pool fueling nuclear growth. Bipartite segregation of pER volume to daughter nuclei at each division cycle modulates nuclear growth kinetics and final size in subsequent embryonic cell divisions. These findings demonstrate that nuclear size regulation is inherently associated with the history of nuclear growth and division.

## RESULTS

### Nuclear growth and division dynamics in early sea urchin development

To visualize nuclear size dynamics *in vivo* during early development, we microinjected unfertilized sea urchin (*Paracentrotus lividus*) eggs with purified GST-GFP-NLS or GST-mcherry-NLS proteins. Nuclear import was apparent soon after fertilization, so that both male and female pro-nuclei became visible ~10-15 minutes after fertilization, shortly before fusing to form the zygote nucleus at the egg center (Tanimoto, et al., 2016) (Fig. 1A, Movie 1). Using spinning disk confocal microscopy, we tracked nuclear surface dynamics in 3D, with a temporal resolution of 2 min over 4-6 hours of embryo development (Fig. 1A and S1A, Movies 2-3). Nuclei in one-cell stage embryos grew continuously until NEB in late prophase. In late anaphase, multiple micronuclei formed around chromosomes (i.e. karyomeres) and eventually fused to form a single new interphase nucleus that resumed growth in the next interphase cycle (Fig. 1A, Movies 1-2).

**Figure 1.**
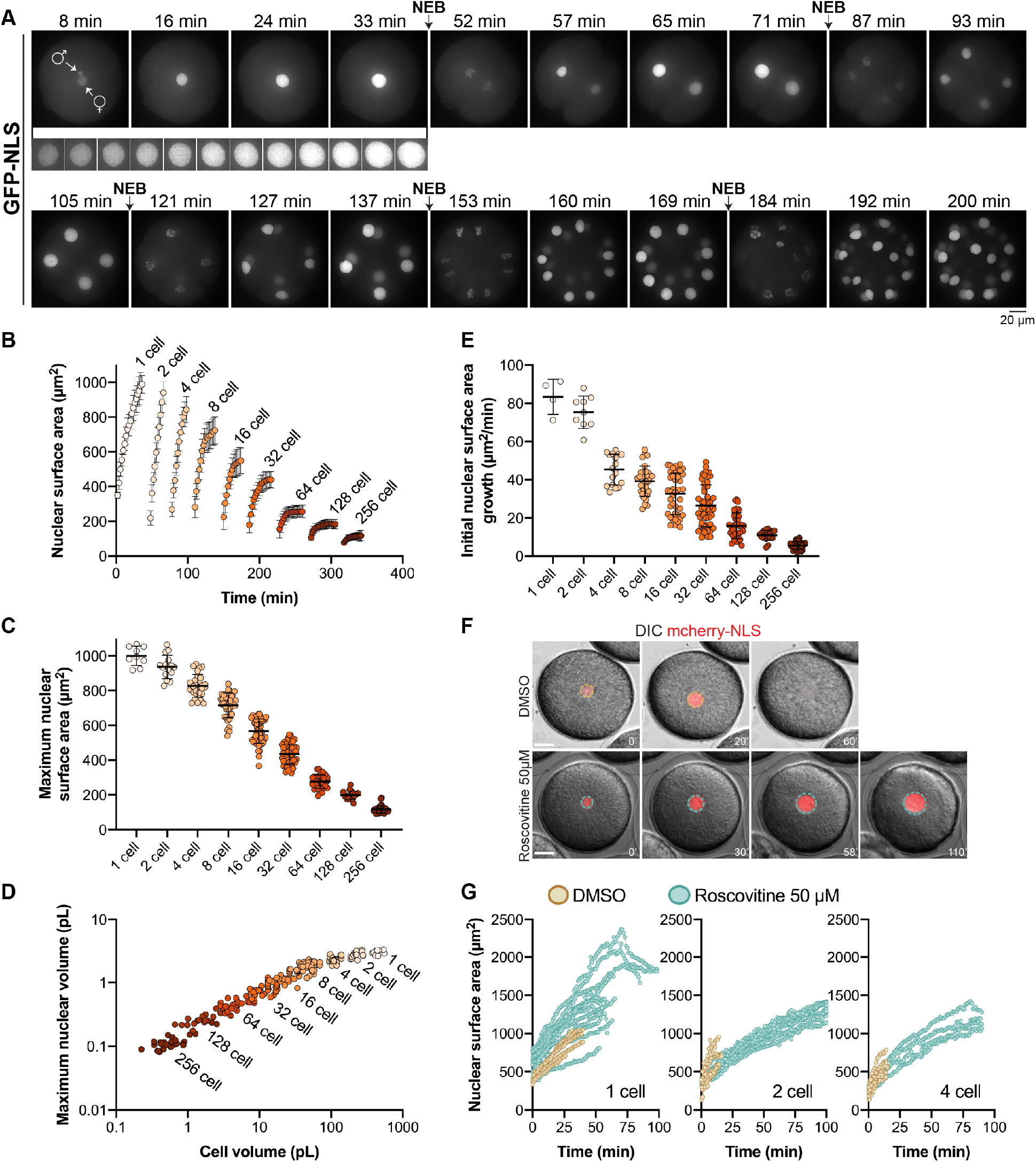
Nuclear size scaling in sea urchin embryos. **(A-E)** Sea urchin eggs were microinjected with GST-GFP-NLS protein prior to fertilization. In some cases, eggs were co-microinjected with mRNA encoding membrane-mCherry and H2B-RFP. Confocal imaging was performed at 1- or 2-minute intervals. Cumulative data from 12 different embryos are shown. Nucleus number: n=9 (1-cell), n=15 (2-cell), n=29 (4-cell), n=41 (8- cell), n=54 (16-cell), n=58 (32-cell), n=34 (64-cell), n=18 (128-cell), n=29 (256-cell). **(A)** Representative maximum intensity z-projections from a time lapse are shown. The male and female pronuclei are indicated in the first image. The inset from 8-33 minutes shows nuclear growth in the 1-cell embryo at 2-minute intervals. NEB refers to nuclear envelope breakdown. Also see Movie 1 and Fig. S1A. **(B)** Maximum nuclear cross-sectional (CS) areas were measured in the GFP-NLS channel. Because the nuclei are roughly spherical (Fig. S1B), we multiplied CS area by 4 to estimate nuclear surface area. Developmental stages were aligned based on when intranuclear GFP-NLS signal was first visible. **(C)** Maximum nuclear surface areas are plotted. **(D)** Individual nuclear and cell volumes are plotted. Nuclear volumes were extrapolated from CS areas (Fig. S1B). Cell volumes were quantified based on membrane-mCherry localized at the plasma membrane (see Fig. S1C). Note that cell volumes were measured for all developmental stages except for 2- and 4- cell embryos where blastomere volumes were calculated as ½ or ¼ of the 1-cell volume, respectively. **(E)** Initial nuclear growth rates were calculated based on the first 3-5 time points of each nuclear growth curve. **(F-G)** Embryos were microinjected with GST-mCherry-NLS protein and were treated with 50 μM roscovitine or an equal volume of DMSO at the 1-cell, 2-cell, or 4-cell stage. Nuclear surface areas were extrapolated from CS areas at one-minute intervals. Note that compared to (A-E), here wide-field imaging was performed with fewer z-planes so size measurements should not be compared between these sets of panels. Control: n=5 (1-cell), n=6 (2-cell), n=7 (4-cell). Roscovitine: n=12 (1-cell), n=10 (2-cell), n=4 (4-cell). Also see Fig. S1H. Error bars represent SD. Scale bars: 20 μm.

Quantification of nuclear surface areas and volumes showed that nuclear growth occurred with nearly no saturation in size in the 1- to 16-cell stages. Nuclei only reached steady state sizes at the 32-cell stage and beyond, with a maximum nuclear size prior to NEB declining steadily from the 1- to 256-cell stage (Fig. 1B-C, Movies 1-3). Maximal nuclear volume decreased ~25-fold from the 1- to 256-cell stage, with a concomitant ~630-fold reduction in blastomere cell volumes (Fig. 1D and S1B-C). However, the correlation between nuclear and cell volumes was highly non-linear, following a power law with an exponent of ~0.37 (Fig. S1D). Importantly, initial nuclear size after NE reassembly was roughly constant from the 2- to 16-cell stages, only decreasing after the 32-cell stage, and therefore did not contribute significantly to the modulation of final nuclear size (Fig S1E). We also noted that interphase length only began to elongate at the 128-cell stage (Fig. S1F), well after the 16-cell stage when nuclei already started reaching steady state sizes. In contrast, the initial nuclear growth rate steadily decreased from the 1-cell stage and beyond, concomitant with reductions in maximum nuclear size (Fig. 1E and S1G), suggesting that surface growth rates play a determinant role in setting final nuclear size.

Accordingly, lengthening the duration of the nuclear growth phase could yield significantly larger nuclei at a fixed cell size. This was evident in embryos treated with roscovitine, an inhibitor of cyclin-dependent kinases that delays NEB (Meijer, et al., 1997) (Fig. 1F and S1H). In roscovitine-treated embryos, nuclei continued to grow well beyond the maximal size of nuclei in controls (Fig. 1G), without increasing their DNA content (Fig. S1I). Delayed nuclei sometimes escaped the cell cycle-block and underwent mitotic disassembly. Others however grew longer, eventually reaching a steady state size with surface areas up to 3.3 times greater than controls (Fig. 1G). This suggests that growing nuclei may be limited in their final size by cell-cycle timing. In summary, nuclei in early cleaving embryos do not typically reach steady-state sizes, but rather exhibit a series of complex growth kinetics that contribute to their size scaling.

### Cell and nuclear sizes are uncoupled

To begin to address mechanisms that dictate developmental nuclear size changes, we set out to thoroughly test the predominant dogma that blastomere cell size dictates nuclear size. We first tested if different volumes of the same cytoplasm could influence nuclear growth and final size following asymmetric divisions. Small vegetal micromeres, for instance, emerge from a programmed asymmetric division at the 8-cell stage (Pierre, et al., 2016) (Fig. 2A). Interestingly, although small micromeres and non-micromere neighbors differed in volume by ~35%, we found that nuclear growth and maximum size were identical (Fig. 2B-C, Movie 4). As a consequence, micromeres exhibited an N/C volume ratio nearly 50% greater than other blastomeres at the same stage (Fig. 2D). Because micromeres could potentially upregulate elements that promote nuclear growth as part of a developmental program, we sought to create ectopic asymmetric divisions at earlier stages. We took advantage of an approach in which microinjected magnetic beads, which recruit endogenous dynein activity, can be cortically positioned to pull on microtubules and associated membrane, generating marked asymmetric cell divisions (Salle, et al., 2019). Remarkably, upon asymmetric division in 1- or 2-cell stage embryos, nuclei also exhibited identical growth kinetics and final size in small and large blastomeres, yielding N/C volume ratios that differed by as much as 25-fold (Fig. 2E-F, Movie 5). In addition, delaying interphase with roscovitine in such asymmetrically divided blastomeres resulted in similar sized nuclei in small and large blastomeres (Fig. S2A-B). These data suggest that cell size has no immediate influence on nuclear growth kinetics or final size and that asymmetric division may spatially pattern the N/C ratio in early embryos.

**Figure 2.**
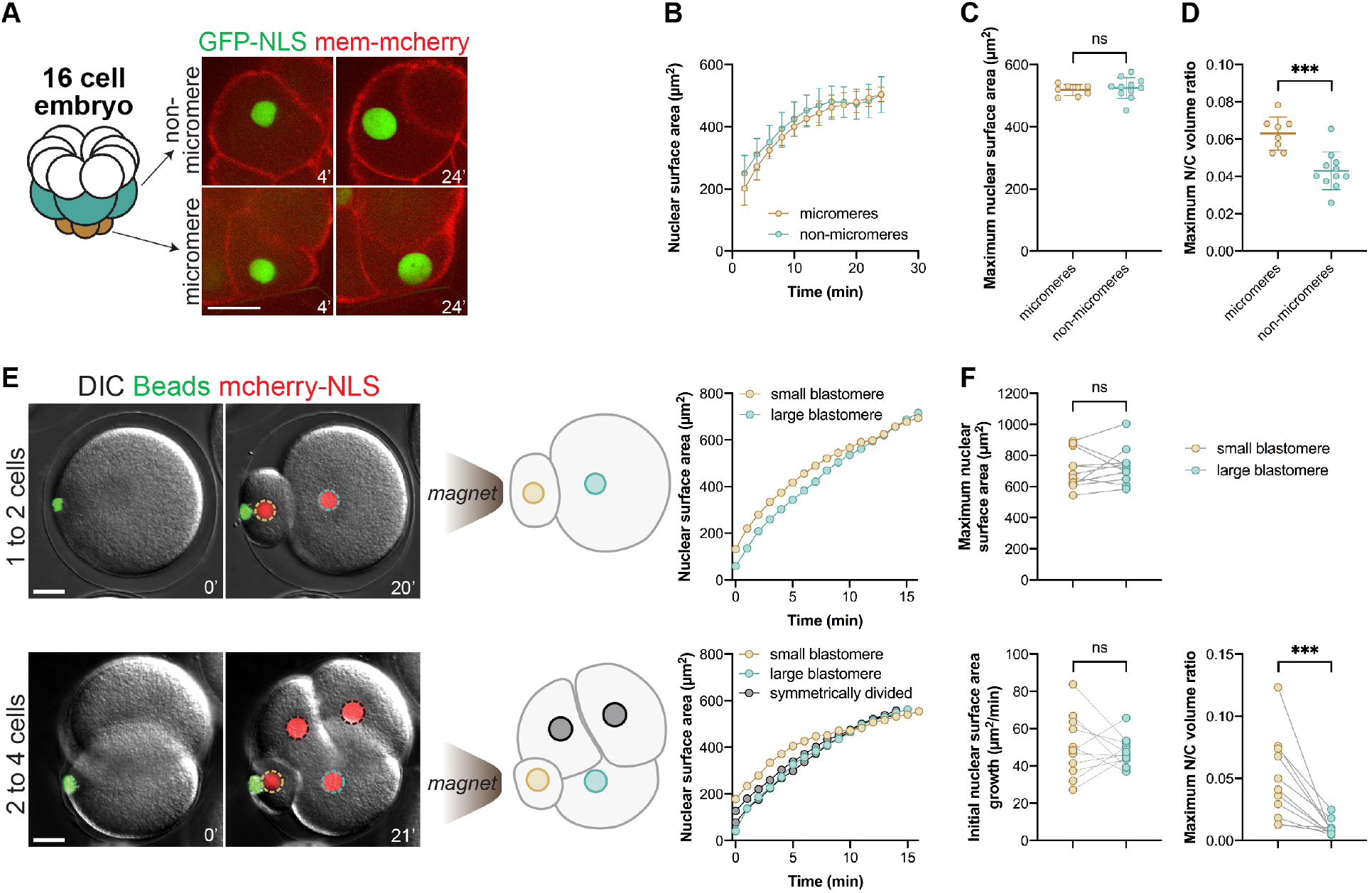
Cell size and nuclear growth are uncoupled in natural and artificial asymmetric cell divisions. **(A-D)** 16-cell stage embryos were analyzed from the experiments described in Fig. 1A-E. Cumulative data from three different embryos are shown: n=8 micromeres, n=11 non-micromeres. Also see Movie 4. **(A)** Representative macromere and micromere. **(B-D)** Nuclear growth curves, maximum nuclear surface areas, and maximum N/C volume ratios are plotted for micromeres and non-micromeres. **(E-F)** Sea urchin embryos were microinjected with GST-mCherry-NLS protein and magnetic beads. An external magnet was used to induce asymmetric divisions at the first or second cleavage. Also see Movie 5. Nuclear surface areas extrapolated from CS areas were quantified at 1-minute intervals based on wide-field imaging. Maximum nuclear surface areas, initial nuclear growth rates, and N/C volume ratios are plotted for 11 small and 11 large blastomeres generated by artificial asymmetric divisions. Wide-field imaging was performed with a limited number of z-planes so these size measurements should not be compared to data obtained from confocal imaging. Error bars represent SD. ***, p<0.005; ns, not significant. Scale bars: 20 μm.

To test the effects of cytoplasmic volume on multiple rounds of nuclear growth and size, we fertilized embryos microinjected with GFP-NLS and then bisected them with a glass needle after zygote nucleus formation and centration. Remarkably, although cut embryos were on average 58 ± 8% the volume of intact controls, nuclear growth kinetics and maximum nuclear sizes were nearly indistinguishable from intact controls across the 4- to 32-cell stages (Fig. 3A-C and S2C-D, Movie 6). We next blocked cytokinesis by treating intact embryos with the aurora kinase inhibitor hesperadin (Argiros, et al., 2012). In these embryos, nuclei number increased in the absence of cell division, generating single-cell multinucleate embryos sharing the same large cytoplasmic volume (Fig. 3D). Growth and final size of individual nuclei in multinucleate embryos were similar to normally dividing controls (Fig. 3D-E). Together these data demonstrate that cytoplasmic volumes have little to no influence on nuclear growth kinetics and final size. Rather, they suggest that nuclear growth and size may be predominantly determined by the history or number of cycles of nuclear assembly, growth, and disassembly through embryo development.

**Figure 3:**
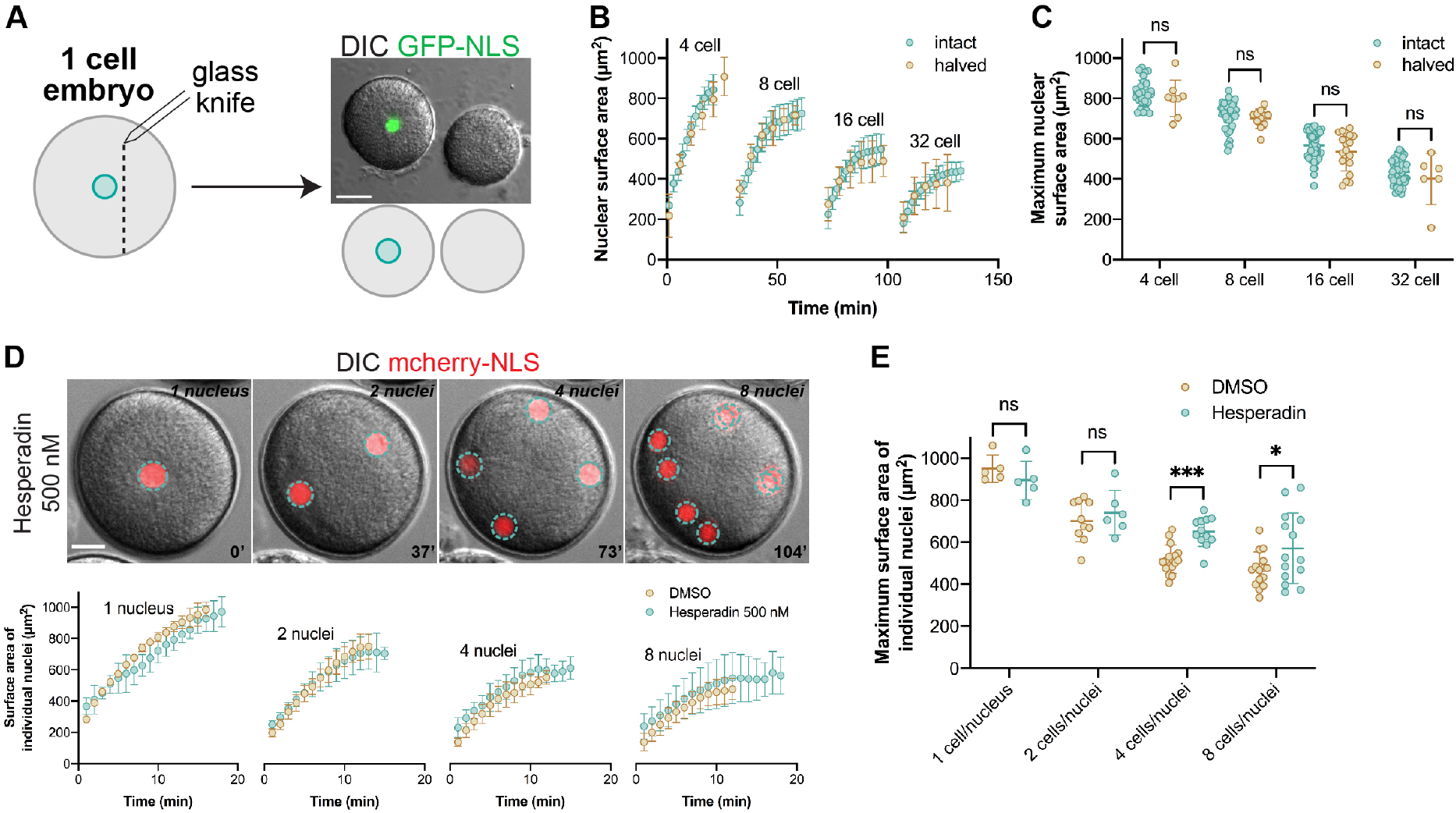
Nuclear growth is unaffected upon increasing the nuclear-to-cytoplasmic ratio. **(A-C)** Sea urchin eggs were microinjected as described in Figure 1, fertilized, bisected with a glass pipet ~30 min post-fertilization, and imaged by confocal at five-minute intervals. Cell volumes in halved embryos were on average 58±8% of intact controls. Nuclear growth curves and maximum nuclear surface areas are plotted for halved and intact embryos based on confocal imaging. Developmental stages were aligned based on when intranuclear GFP-NLS signal was first visible. Intact embryo data are the same shown in Fig. 1B-C. Cumulative data from two different halved embryos are shown: n=8 (4-cell), n=11 (8-cell), n=18 (16-cell), n=6 (32-cell). Maximum N/C volume ratios in halved embryos were on average 1.7±0.5 fold greater than in intact controls. **(D-E)** One-cell embryos were microinjected with GST-mCherry-NLS protein and were treated with 500 nM hesperadin or an equal volume of DMSO. Nuclear surface areas extrapolated from CS areas were quantified for individual nuclei at one-minute intervals based on wide-field imaging. Nuclear growth curves and maximum nuclear surface areas for individual nuclei are plotted. DMSO: n=5 (1-cell), n=10 (2-cell), n=16 (4-cell), n=15 (8-cell). Hesperadin: n=5 (1-nucleus), n=6 (2-nuclei), n=12 (4-nuclei), n=14 (8-nuclei). Wide-field imaging was performed with a limited number of z-planes so these size measurements should not be compared to data obtained from confocal imaging. In hesperadin-treated embryos, we sometimes noted nuclear fusion. Error bars represent SD. ***, p<0.005; *, p<0.05; ns, not significant. Scale bars: 20 μm.

### The perinuclear ER fuels nuclear growth

The saturating behavior of nuclear growth curves beyond the 16-cell stage and in a subset of roscovitine-delayed nuclei indicated the existence of putative limiting components that fuel nuclear growth (Fig. 1B and 1G). However, if size-limiting components were freely diffusible in the cytoplasm, nuclear size would have been expected to be sensitive to cell size (Goehring and Hyman, 2012). Given the lack of influence of cytoplasmic volume on nuclear growth and size in our system, we thus searched for limiting components that would be restricted to a local region around nuclei, rather than being distributed evenly throughout the cytoplasm.

Nuclear growth has been previously shown to depend on nuclear import (Newport, et al., 1990). Import could potentially be titrated in a local manner at the NE during development, for instance by a scaled-regulation of the activity or density of NPCs. We thus measured nuclear import kinetics over early development (Figs. 1A, S3A-C). While nuclear growth and size generally correlated with import rates, which steadily decreased over development in accordance with *Xenopus* studies (Levy and Heald, 2010), we noted that in very early stages a plateau was reached such that further increase in import rates did not lead to faster nuclear growth (Fig. S3C-E). Nuclear import rates were also not significantly altered in halved embryos and were similar in 16-cell stage micromeres and non-micromeres (Fig. S3F-G). However, in 2- and 4-cell stage embryos forced to divide asymmetrically with magnetic beads, nuclear import was greater in large blastomeres even though nuclear growth was similar in small and large blastomeres (Fig. 2E-F and S3H). Although these data do not definitively disprove a role for import in nuclear growth, they suggest that growth and import rates can be uncoupled, indicating that import may not serve as the prime local factor limiting and scaling nuclear growth.

Another essential cellular component that has been suggested to affect nuclear morphology, NE assembly, and nuclear remodeling is the ER. Indeed, the ER constitutes a large reservoir of membranes and lipids that may contribute to NE remodeling and expansion (Anderson and Hetzer, 2008; Anderson and Hetzer, 2007). In agreement with its potential role as a local nuclear growth regulator, immunostaining revealed a massive perinuclear accumulation of ER in 1-cell stage embryos (Fig. 4A). This perinuclear ER (pER) spanned a roughly toroidal region ranging from 3-10 μm wide (~5 μm on average). This accumulation appeared to be progressive, initiating early after fertilization around the centering male pronucleus in agreement with previous reports (Terasaki and Jaffe, 1991) (Fig. 4A, Movie 7).

**Figure 4.**
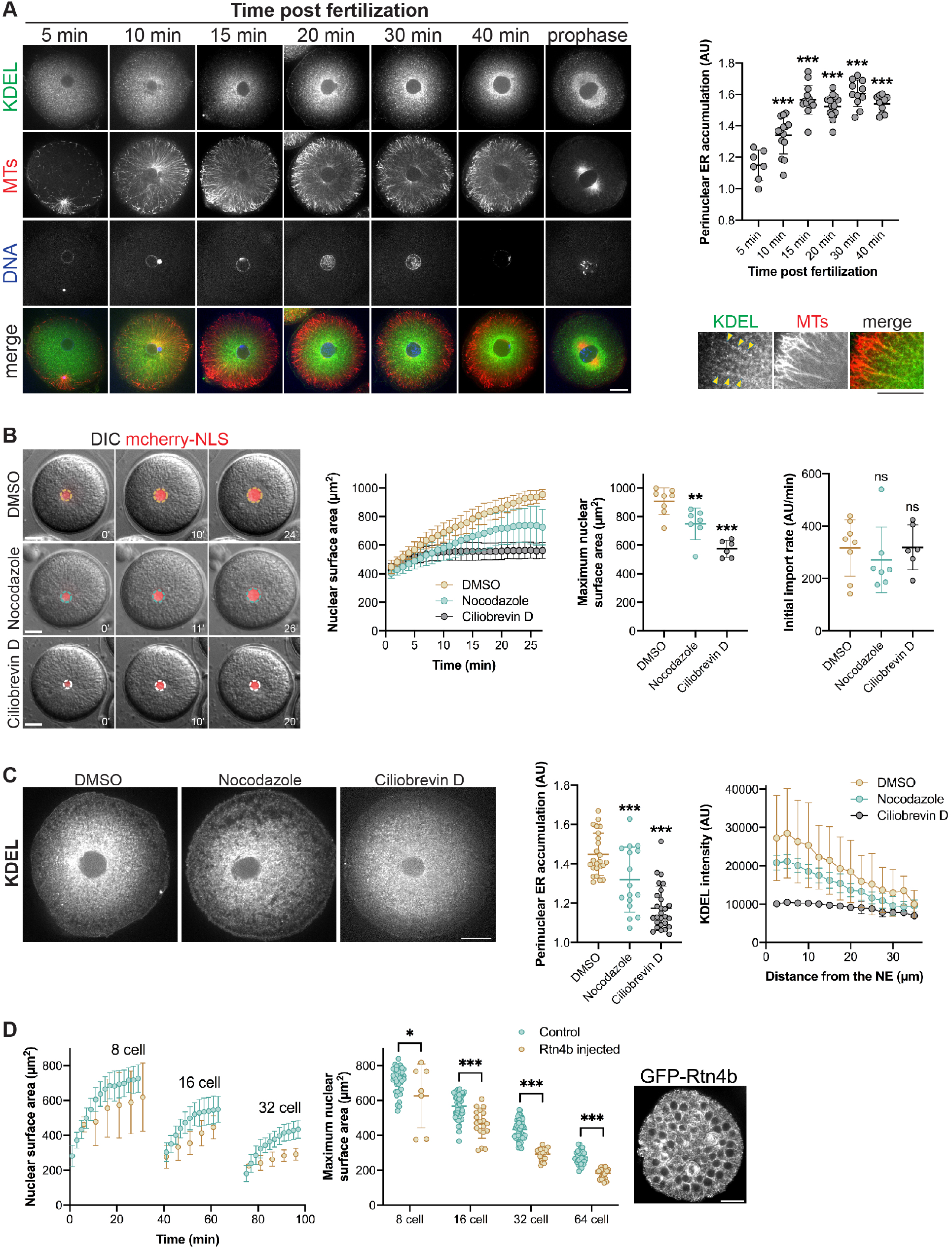
Disrupting perinuclear ER accumulation reduces nuclear growth. **(A)** One-cell sea urchin embryos were fixed at different times after fertilization and immunostained with anti-KDEL and anti-tubulin antibodies. To quantify perinuclear ER accumulation, the mean KDEL intensity within a concentric ring measuring 15 μm from the NE was divided by the mean KDEL intensity of the whole embryo excluding the nucleus. n=7 (5 min), n=14 (10 min), n=14 (15 min), n=16 (20 min), n=11 (30 min), n=10 (40 min). The inset shows co-localization of MTs and ER membrane. The arrowheads mark KDEL puncta on MTs. **(B)** One-cell sea urchin embryos microinjected with GST-mCherry-NLS were treated after aster centration with 20 μM nocodazole to depolymerize microtubules (n=7), 50 μM ciliobrevin D to inhibit dynein (n=6), or DMSO (n=8) as a control. Nuclear sizes were quantified at one-minute intervals based on wide-field imaging starting with the first appearance of intranuclear mCherry-NLS signal. Nuclear growth curves, maximum nuclear surface areas, and initial nuclear import rates are plotted. See Fig. S3 and Methods for details on how import rates were calculated. Wide-field imaging was performed with a limited number of z-planes so these size measurements should not be compared to data obtained from confocal imaging. **(C)** Twenty minutes after fertilization, embryos were treated with 20 μM nocodazole, 50 μM ciliobrevin D, or DMSO as a control. Embryos were fixed 10 minutes later and immunostained with an anti-KDEL antibody. Also see Fig. S4A. Note that these KDEL images are the same shown in Fig. S4A for 10 minutes after drug addition. Perinuclear ER accumulation was quantified as in (A) for 16-27 nuclei per condition. To measure the KDEL distribution relative to the NE, KDEL intensity was quantified within concentric rings expanding away from the NE (see Methods) for 14-15 nuclei per condition. **(D)** Sea urchin eggs were microinjected with GST-mCherry-NLS protein and mRNA encoding GFP-Rtn4b prior to fertilization. A representative blastula stage embryo is shown to the right. Time lapse confocal imaging was performed at 5-minute intervals for two different GFP-Rtn4b microinjected embryos: n=7 (8-cell), n=19 (16-cell), n=20 (32-cell), n=26 (64-cell). Nuclear growth curves and maximum nuclear surface areas are plotted. Control data are the same shown in Figure 1B-C. Error bars represent SD. ***, p<0.005; **, p<0.01; *, p<0.05; ns, not significant. Scale bars: 20 μm.

To begin to test if pER might regulate nuclear growth, we first manipulated the microtubule (MT) cytoskeleton. Indeed, as previously described in *Xenopus* extracts (Wang, et al., 2013), the ER co-aligns with MTs in large interphase asters, suggesting that pER accumulation could rely on MT minus-end directed transport powered by dynein (Fig. 4A). Accordingly, dynein inhibition in single-cell embryos largely dispersed the pER within minutes and completely abrogated nuclear growth. Acute MT depolymerization, however, only partially disrupted ER structure with a resultant mild reduction in nuclear growth (Fig. 4B-C and S4A). We suspect that dynein inhibition had a more dramatic effect because the activity of MT plus-end directed motors or MT polymerization would tend to disperse the ER away from the nucleus (Wang, et al., 2013). Interestingly, in spite of dynein inhibition and MT depolymerization reducing nuclear growth, the initial nuclear import rate was unaffected, providing further evidence that import alone cannot support and scale nuclear growth (Fig. 4B).

To more directly test the effect of ER morphology on nuclear growth, we ectopically expressed Reticulon 4b (Rtn4b), an ER tubule-shaping protein that is known to disrupt pER sheets (Shibata, et al., 2010). It has been proposed that a tug-of-war exists between ER and nuclear membranes such that a more tubulated ER restricts membrane flow to the nucleus (Anderson and Hetzer, 2008). Consistent with results in *Xenopus* embryos (Jevtic and Levy, 2015), Rtn4b expression reduced nuclear growth and size in sea urchin embryos, with stronger effects observed later in development likely due to increased accumulation of Rtn4b protein expressed from microinjected mRNA (Fig. 4D). We conclude that perinuclear accumulated ER could serve as a local membrane source fueling, and potentially limiting, nuclear surface growth.

### Size scaling of perinuclear ER can account for nuclear growth and size scaling during embryogenesis

Given that altering ER structure and perinuclear accumulation affected nuclear growth, we wondered if pER amount may change over the course of normal development, potentially contributing to nuclear size scaling. Strikingly, imaging of immunostained pER at different developmental stages revealed a progressive reduction in pER during the course of development, correlating with developmental reductions in nuclear size (Fig. 5A). Volume quantification of pER through automated segmentation in 3D (Fig. S4B) showed that pER volume is reduced by a factor of ~2.2 at each division cycle (Fig. 5B-C). This reduction was confirmed by live imaging of KDEL-labeled ER at the 8- to 32-cell stages (Fig. S4C). Live-imaging of embryos injected with DiI-dye that labels the ER (Terasaki and Jaffe, 1991) further confirmed this ~2-fold reduction at each cell division, and revealed that it occurred mostly as a consequence of mitosis, with only minor pER depletion during interphase (Fig. 5D and Movie 7). At the onset of mitosis, the local pER pool remained in the vicinity of the disassembling nucleus and became associated with mitotic asters that split it into two independent pools at anaphase. Importantly, this bipartite segregation was conserved in asymmetrically dividing micromeres suggesting it was not influenced by cell size (Fig. 5B). These data demonstrate that after cell division, growth of newly formed nuclei may be fueled by an amount of local pER material, roughly half that associated with nuclei in the previous cycle and independent of cytoplasmic volume.

**Figure 5:**
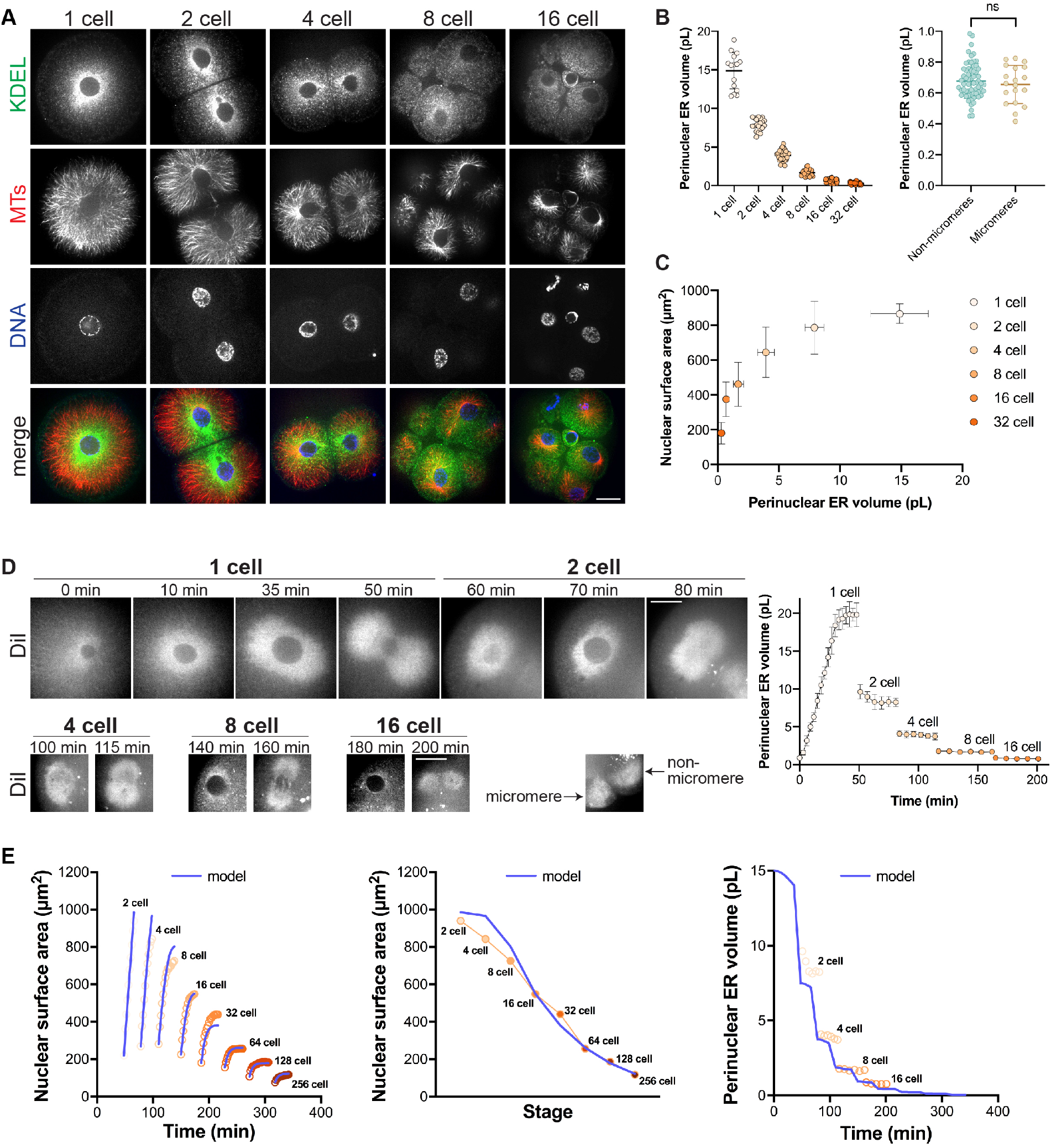
Perinuclear ER volume halves during sequential embryonic divisions. **(A-C)** Sea urchin embryos at different developmental stages were fixed and immunostained with anti-KDEL and anti-tubulin antibodies. **(A)** Representative images are shown. **(B)** Perinuclear ER volume was quantified from confocal z-stacks (see Fig. S4B and Methods). Cumulative data are shown for fixed KDEL-stained embryos and for live embryos microinjected with mRNA encoding mCherry-KDEL (see Fig. S4C). n=15 cells (1-cell), n=20 cells from 10 embryos (2-cell), n=35 cells from 9 embryos (4-cell), n=23 cells from 3 embryos (8-cell), n=103 cells from 12 embryos (16-cell), n=34 cells from 6 embryos (32-cell), n=63 cells (non-micromeres), n=19 cells (micromeres). **(C)** Nuclear surface areas were measured for the same cells described in (B). Average nuclear surface areas are plotted as a function of average pER volumes for different stages. **(D)** Embryos stained with DiI were imaged by time-lapse confocal microscopy. Perinuclear ER volume was quantified from confocal z-stacks (see Fig. S4B and Methods). Cumulative data from 6 embryos are shown. n=6 (1-cell), n=11 (2-cell), n=14 (4-cell), n=20 (8-cell), n=6 (16-cell). **(E)** Model predictions are overlaid on experimental data for nuclear growth curves, final nuclear sizes, and pER volumes. The modeling output is based on the parameter values reported in STAR Methods. Error bars represent SD. ns, not significant. Scale bars: 20 μm.

To test if the bipartite segregation of pER material at each division could account for the modulations of nuclear growth kinetics and size scaling, we developed a minimal kinetic model for nuclear growth. This model posits a transfer of pER material amount to the nuclear surface, occurring at a rate *k*_0_, independent of embryonic progression or cell size. Amount transfer best represents membrane transfer, as this process is not expected to be limited by diffusion. Because pER was not fully depleted in each interphase, we posited that only a fraction λ of the pER is available for transfer to the nucleus. Calling 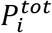 the volume of pER at stage *i*, the available fraction is thus 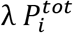. However, the pER has a complex topology and the relationship between pER volume and available surface material for NE expansion is not necessarily linear. Therefore, we introduced a phenomenological scaling parameter *α* so that the total available surface material provided by a transfer from the pER at stage *i* is:

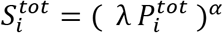

Experimental traces of nuclear growth were indicative of a saturated Michaelis-Menten type kinetics (Fig. 1B). Assuming such kinetics for nuclear growth required the introduction of one saturation parameter, *S*^*sat*^. The time evolution of nucleus size at each stage *i*, *N*_*i*_(t) thus reads:

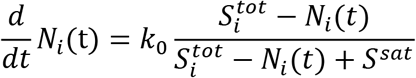

Finally, the equal partitioning of pER volume at each division yields:

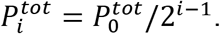

*N*_*i*_(t) was solved analytically and the parameters were fitted to the experimental nuclear growth curves from the 2- to 256-cell stages (see Star Methods). Remarkably, this simple model could accurately account for all details of the series of nuclear growth kinetics, as well as final nuclear size scaling in early embryos (Fig. 5E). Importantly, assuming no dependence on pER for the available surface material (*α* = 0) yielded self-similar growth curves through embryo development, with no reduction in nuclear size. Conversely, taking a linear relationship between pER volume and available surface material (*α* = 1) yielded a rapid depletion of available material preventing any further nuclear growth after the 8-cell stage (Fig. S4D). The best fitting value for the exponent *α* was 0.54, smaller than, yet close to, a value of 2/3 that would correspond to a surface-to-volume relation converting the pER volume to an available surface material for nuclear surface growth. Thus, this minimal model demonstrates that the progressive reduction in pER material can explain the rich kinetics of nuclear growth during development, with a single fixed reaction rate.

### Validation of the model in a vertebrate system

We next tested if our findings are conserved between invertebrate and vertebrate embryos. We first imaged pER in *Xenopus* embryos. As in sea urchin embryos, immunostaining in *Xenopus* blastomeres revealed a reduction in pER volume with each division and a scaling between nuclear size and pER volume (Fig. 6A-B), resembling that obtained in sea urchin embryos (Fig. 5A-C). Thus, the gradual reduction of pER amount as a result of successive cell divisions may be a general feature of developing invertebrate and vertebrate embryos.

**Figure 6.**
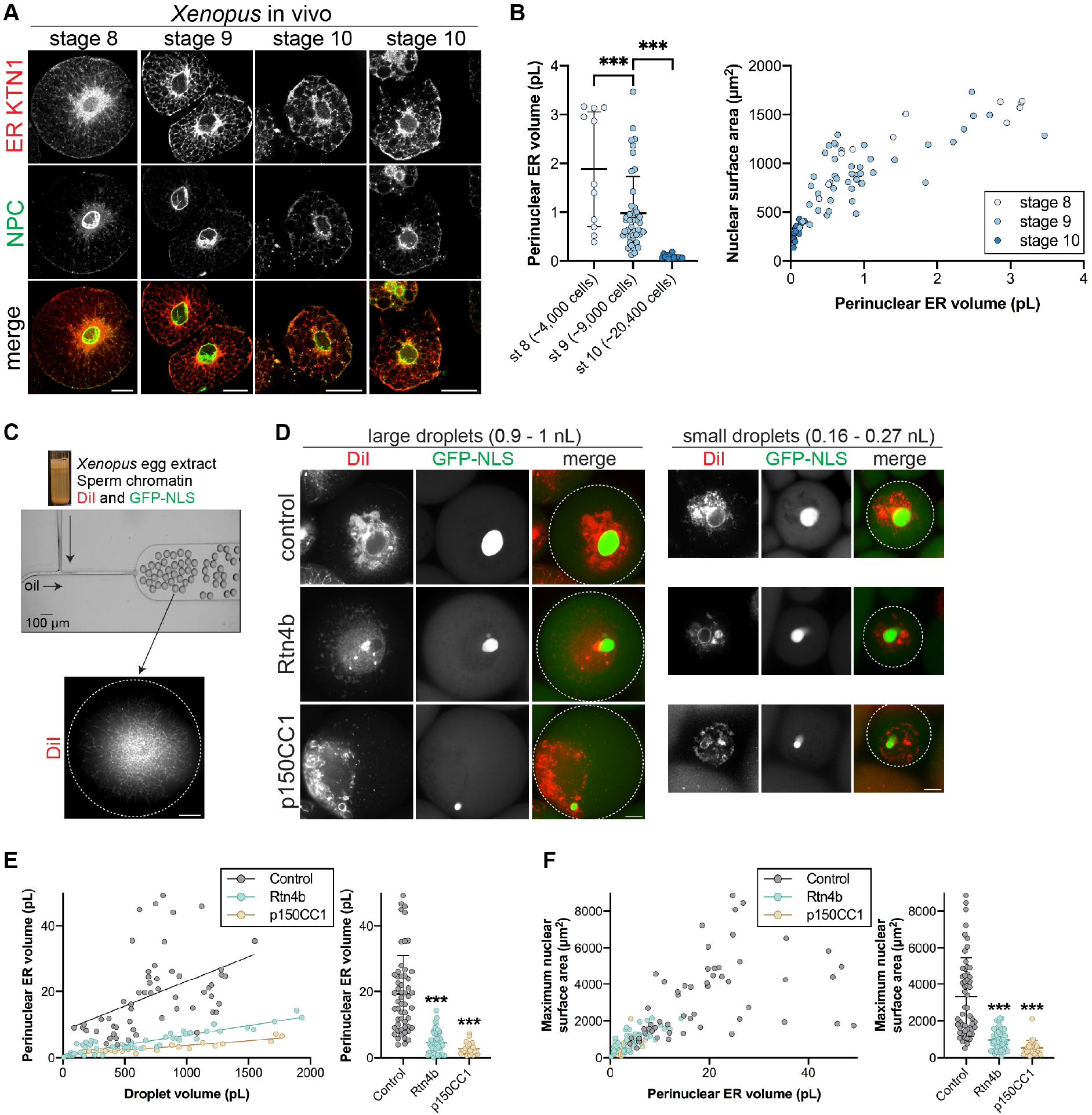
Perinuclear ER volume scales nuclear size *in vitro*. **(A-B)** Blastomeres from different stage *Xenopus laevis* embryos were dissociated, fixed, and stained with antibodies against perinuclear ER sheet marker KTN1 and the NPC (mAb414). **(A)** Representative confocal images are shown. **(B)** Perinuclear ER volume was quantified from confocal z-stacks (see Fig. S4B and Methods). Nuclear surface area was extrapolated from CS area. Cumulative data are shown from three different batches of embryos. n=11 (stage 8), n=46 (stage 9), n=14 (stage 10). **(C-F)** Nucleus and ER formation were induced in fractionated interphase *X. laevis* egg extract supplemented with membrane dye DiI and GST-GFP-NLS protein. Extract and nuclei were encapsulated in droplets of differing volumes using microfluidic devices. Where indicated, extracts were supplemented with 67 nM recombinant Rtn4b protein or 3.2 μM recombinant p150CC1 dynein inhibitor. Confocal z-stacks were acquired after a 3-hour incubation at 16°C. n=55 (control), n=52 (Rtn4b), n=27 (p150CC1). **(C)** The experimental approach. The image of the microfluidic device was adapted with permission from (Hazel, et al., 2013). **(D)** Representative images of different sized droplets. Imaging of the droplet periphery verified that Rtn4b addition induced a more tubulated cortical ER (data not shown). The droplet boundary is outlined in white in the merged images. **(E)** Perinuclear ER volume was quantified from DiI images for different size droplets (see Fig. S4B and Methods). Previous studies have established that DiI stains ER membranes in *Xenopus* extract (Dreier and Rapoport, 2000). **(F)** Nuclear CS area was quantified from GFP-NLS images for different size droplets and extrapolated to surface area (see Methods). For individual droplets, nuclear surface area was plotted as a function of the perinuclear volume measured in (E). Error bars represent SD. ***, p<0.005. Scale bars: 20 μm.

Given the opacity of *Xenopus* embryos, we next assayed the requirement of pER for nuclear growth using *Xenopus* cell-free extracts that faithfully reconstitute nuclear and ER biology (Chen and Levy, 2018; Dreier and Rapoport, 2000). Egg extracts were supplemented with sperm chromatin, DiI-dye, and GFP-NLS, activated with calcium, and then rapidly encapsulated in droplets of different sizes using microfluidic devices (Hazel, et al., 2013) (Fig. 6C-D). In both large and small droplets, nuclei grew to reach a final steady state size, typically within 120 min after calcium activation (Fig. S5A). Because nuclei are initially the same size, the steady state size serves as a proxy for nuclear growth rates. As *in vivo*, the ER also accumulated around nuclei surfaces (Fig. 6D). Importantly, inhibiting dynein, by addition of the p150-CC1 dominant negative fragment, prevented pER accumulation and strongly reduced the final size of nuclei (Fig. 6D-F and S5B). Addition of recombinant Rtn4b reduced pER sheet formation and final nuclear size (Fig. 6D-F, S5B-D). Thus, in agreement with data from live urchin embryos, pER accumulation appears to be required for nuclear growth. Furthermore, by varying droplet size we modulated the total amount of encapsulated ER, which resulted in larger pER volumes in larger droplets. Remarkably, droplets containing larger volumes of pER produced nuclei with larger final sizes (Fig. 6D-F and S5B). Because pER is equally divided between cells in vivo during normal divisions, our finding that in vitro pER volume correlates with nuclear size reproduces the scaling seen in vivo and in our model. Thus, pER membrane amount may represent a generic predictor of nuclear growth rates and final size in early embryos, independently of cell size.

## DISCUSSION

### Nuclear size scaling during development is independent of cell size

Based on live imaging of transparent sea urchin embryos, we here report on the dynamics of nuclear size throughout early development of an intact embryo. These data unravel essential details of nuclei growth kinetics and import rates, unattainable in previous studies that were often based on measurements at a limited number of developmental time points or in fixed embryos (Jevtic and Levy, 2015; Levy and Heald, 2010). One significant finding is that nuclei continuously grow during interphase, only reaching saturating sizes beyond the 32-cell stage. This renders final nuclear sizes heavily dependent on growth rates as well as interphase duration, important considerations beyond the notion of mere “size-regulating” elements. Furthermore, through multiple independent experiments, we largely exclude any influence of cell size on nuclear size. Thus, these data may call for a reinterpretation of the established correlation between cytoplasmic volumes and nuclear size. Although we cannot rule out species-specific strategies, we speculate that this correlation in other systems could be explained by the concomitancy of cell and nuclear growth (Walters, et al., 2019; Jorgensen, et al., 2007; Neumann and Nurse, 2007), tissue and nuclear growth (Windner, et al., 2019), or the number of cell and nuclear divisions.

The lack of direct influence of cell size on nuclear size impacts N/C ratio values during development, with potential implications for zygotic genome activation (ZGA) and fate specification in early embryos (Jevtic and Levy, 2015). First, our measurements show that the N/C volume ratio increases over the course of early development, in both urchins and frogs, with higher values reached much earlier in urchins (Fig. S5E). Because ZGA is already detectable soon after fertilization in urchins (Poccia, et al., 1985; Nemer, 1963) but only becomes prevalent at the ~4000-cell stage in frogs, this raises an interesting hypothesis that ZGA may be triggered upon crossing a conserved N/C threshold in different species. Second, because nuclear size is independent of cell size, asymmetric division emerges as a dominant strategy to spatially regulate the N/C ratio. Accordingly, recent studies in *Xenopus* suggest that ZGA may be spatially patterned along the animal-vegetal axis, correlating with a gradient in cell sizes generated through rounds of asymmetric divisions along this axis (Chen, et al., 2019). In many invertebrate and vertebrate embryos, asymmetric divisions in size are also often linked to cell fate specification and modulations of cell cycle length (Hawkins and Garriga, 1998). For instance, sea urchin micromeres formed by asymmetric divisions at the 8-cell stage typically slow down their cell cycle length and relocate fate markers such as β-catenin in their nuclei (Gilbert, 2010). Addressing if such changes are direct consequences of altered N/C ratios caused by the reduction of cytoplasmic volume at a fixed nuclear size, or solely part of a developmental program, is an exciting future research avenue.

### Role of the perinuclear ER in nuclear growth and size control

Because cell size may not serve as a prime determinant of nuclear growth and size in embryos, we suggest that nuclear growth is not limited by soluble components evenly distributed throughout the cytosol. Rather, our data support the idea that dynein-mediated accumulation of pER material is essential to drive nuclear surface growth, providing *in vivo* validation of a previous model proposed from frog extracts (Hara and Merten, 2015). Consistent with this model, we found that reducing cell volume had little effect on nuclear volume as long as the ER membranes remain associated with the nucleus. Conversely, in other systems clustered nuclei tend to grow less because they presumably have to share the same local perinuclear membrane material (Windner, et al., 2019; Hara and Merten, 2015; Neumann and Nurse, 2007; Gurdon, 1976). In addition, modulating ER amount by stratification of the cytoplasm demonstrated a role for the ER in male pronuclear growth and maturation at fertilization (Longo, 1976). Thus, the local pool of available ER or other membranes may be one conserved element for fueling nuclear membrane growth (Walters, et al., 2019; Hara and Merten, 2015; Golden, et al., 2009).

Although independent of cell size, such a model is still consistent with limiting components scaling nuclear size during development because limiting local perinuclear material appears to be serially distributed between greater numbers of nuclei over development. Our data revealed that pER volume was reduced by a factor of ~2 at each cell division, likely though ER membrane association with dividing mitotic asters. As demonstrated by our minimal mathematical model, this bipartite segregation of the pER pool may be sufficient to account for the full range of developmental modulations in nuclear growth kinetics and final size. Although not needed in this minimal model, it is likely that other effects could influence pER amount in successive divisions, including gradual changes in membrane synthesis, MT/dynein transport efficiency (Olenick and Holzbaur, 2019; Lacroix, et al., 2018), as well as expression and/or relocalization of ER shaping proteins (Schwarz and Blower, 2016).

Finally, our data raise important questions on the mechanobiological mechanisms that control and/or limit nuclear growth. Although the pER appears to be required for nuclear growth, nuclear import has also been implicated in nuclear size control (Brownlee and Heald, 2019; Levy and Heald, 2010). Results from a genome-wide screen in yeast suggest that both mechanisms may control nuclear size in concert (Kume, et al., 2017). In our data set, import rates generally scaled with nuclear surface growth rates, although in several instances the two parameters could be uncoupled. In light of these observations, we suggest that import is required, but not necessarily limiting, for nuclear growth. Akin to the growth of pressurized thin shells, like walled cells, which requires a turgid cytoplasm but is limited by membrane and cell wall insertion (Davi, et al., 2018), we speculate that nuclear import could contribute to pressurizing the nucleoplasm but that membrane addition from the pER may confer viscoelastic properties that allow for net nuclear surface expansion. Further work addressing how the physical properties of nuclei are modulated throughout development will be required to build mechanistic models for the control of nuclear growth and size.

## Abbreviations

NE: nuclear envelope
NPC: nuclear pore complex
ER: endoplasmic reticulum
pER: perinuclear endoplasmic reticulum
NLS: nuclear localization signal
CS: cross-sectional
N/C: nuclear-to-cytoplasmic
ZGA: zygotic genome activation
NEB: nuclear envelope breakdown
MT: microtubule

## ACKNOWLEDGEMENTS

We thank Matthew Dilsaver for purifying recombinant Rtn4b protein and Amanda Johnson for validating its ER localization, Abdullah Bashar Sami, Miroslav Tomschik, and Jay Gatlin for providing recombinant p150CC1, and Eloise Fadial for fabricating microfluidic devices. Research in the Levy lab is funded by the National Institutes of Health/National Institute of General Medical Sciences (R01GM113028 and P20GM103432) and the American Cancer Society (RSG-15-035-01-DDC). Research in the Minc lab is funded by the Centre National de la Recherche Scientifique and the European Research Council (CoG Forcaster no. 647073). SD acknowledges funding from the CNRS Momentum. We acknowledge the ImagoSeine core facility of the Institut Jacques Monod, member of Infrastructure en Biologie Santé et Agronomie and France-BioImaging (ANR-10-INBS-04) infrastructures.

## AUTHOR CONTRIBUTIONS

Conceptualization, RNM, JS, NM, DLL; Methodology, KN, JO; Modeling, SD and NM; Investigation, RNM, JS, SD, NM, DLL; Writing – Original Draft, RNM, JS, NM, DLL; Writing – Review & Editing, RNM, JS, SD, KN, JO, NM, DLL; Funding Acquisition, NM, DLL; Supervision, NM, DLL

## STAR METHODS

### CONTACT FOR REAGENT AND RESOURCE SHARING

Further information and requests for resources and reagents should be directed to and will be fulfilled by the lead contact, Daniel Levy.

### EXPERIMENTAL MODEL AND SUBJECT DETAILS

#### Sea urchin maintenance and gametes collection

Purple sea urchins (*P. lividus*) were obtained from the Roscoff Marine station (France) and kept at 16°C in an aquarium for several weeks in artificial seawater (Reef Crystals; Instant Ocean). Gametes were collected by intracoelomic injection of 0.5 M KCl. Sperm was collected dry and kept at 4°C for 1 week. Eggs were rinsed twice, kept at 16°C, and used on the day of collection.

#### *Xenopus* embryos

*X. laevis* embryos were obtained as previously described (Sive, et al., 2000). Freshly laid *X. laevis* eggs were in vitro fertilized with crushed *X. laevis* testes. Embryos were dejellied in 3% cysteine (w/v) pH 7.8 dissolved in 1/3x MMR (1× MMR: 0.1 mM EDTA, 0.1 M NaCl, 2 mM KCl, 2 mM CaCl_2_, 1 mM MgCl_2_, 5 mM HEPES pH 7.8). Embryos were developed in 1/3× MMR, staged, and arrested in late interphase with 0.15 mg/mL cycloheximide for 60 mins unless otherwise indicated. All *Xenopus* procedures and studies were conducted in compliance with the US Department of Health and Human Services Guide for the Care and Use of Laboratory Animals. Protocols were approved by the University of Wyoming Institutional Animal Care and Use Committee (Assurance #A-3216-01).

#### METHOD DETAILS

##### Sea urchin microscopy

Microinjections and magnetic tweezers manipulations were performed on a wide-field fluorescence microscope (TI-Eclipse; Nikon) equipped with a complementary metal oxide-semiconductor camera (Orca-flash4.0LT; Hamamatsu). Samples were imaged on the injection setup with a 20× dry objective (NA, 0.75; Apo; Nikon) and a 1.5× magnifier (final pixel size, 0.217 μm); or alternatively on a separate fluorescent microscope (Leica DMI 6000B) equipped with the same camera and a 20× dry objective (NA, 0.70; PLAN Apo; Leica) (final pixel size, 0.460 μm). Microscopes were operated with Micro-Manager (Open Imaging). Live imaging was carried out in artificial seawater at a stabilized room temperature (18–20°C). Time-lapse confocal imaging was performed using a spinning-disk microscope (TI-Eclipse, Nikon) equipped with a Yokogawa CSU-X1FW spinning head and an EM-CCD camera (Hamamatsu) using a 60× water immersion objective (NA, 1.2; PLAN Apo; Nikon). Confocal imaging of fixed samples was performed on a Zeiss LSM780 with a 63× water immersion objective (NA, 1.4; C-Apo; Zeiss). Time lapses were generally acquired at 1-5 minute intervals with z distance step sizes of 1 μm.

##### Sea urchin embryo microinjections and manipulations

###### Microinjections

The jelly coat of unfertilized eggs was removed by passing them three times through an 80 μm Nitex mesh (Genesee Scientific) to facilitate egg adhesion on protamine-coated glass-bottom dishes (MatTek Corporation). Unfertilized eggs were transferred to protamine-coated glass-bottom dishes for a maximum time of 15 min before microinjection and fertilization. Microinjections were performed using a FemtoJet 4 and Injectman 4 micromanipulator (Eppendorf). Injection pipettes were prepared from siliconized (Sigmacote; Sigma-Aldrich) borosilicate glass capillaries (1 mm diameter). Glass capillaries were pulled with a needle puller (P-1000; Sutter Instruments) and ground with a 30° angle on a diamond grinder (EG-40; Narishige) to obtain a 5-10 μm aperture. Injection pipettes were back-loaded with 2 μl before each experiment and were not reused. Injection volumes were generally less than 5% of the egg volume, ~2-5 pL. GST-GFP-NLS and GST-mCherry-NLS proteins were diluted to ~2.5 mg/ml in PBS before injecting. Injecting undiluted GST-GFP-NLS did not affect developmental progression, suggesting that these protein concentrations are not detrimental to the embryo. RNA was diluted to ~50 ng/μl in PBS before injecting. These concentrations were empirically selected to optimize imaging while not affecting developmental progression. To label ER in live embryos, DiIC_18_(3) powder (Molecular probes) was mixed with soybean oil and incubated for 12h in the dark at room temperature. Oil was cleared by centrifugation on a benchtop centrifuge to remove excess crystals. An oil droplet (approximately 5-10 μm in diameter) was injected in eggs 15-20min before fertilization to allow the dye to diffuse into ER membranes.

###### Chemical inhibition

For drug treatments, 100× stock aliquots were prepared in DMSO. Nocodazole (Sigma-Aldrich) was used at a final concentration of 20 μM. Ciliobrevin D (EMD Millipore) was used at a final concentration of 50 μM. Roscovitine (Sigma-Aldrich) was used at a final concentration of 50 μM. Hesperadin (SelleckChem) was used at a final concentration of 500 nM. To image DNA, Hoechst 33342 (Sigma-Aldrich) was used at a final concentration of 10 μg/ml. As these drugs are cell-permeable, they were added directly to the seawater in which the embryos were cultured.

###### Egg bisection

After microinjection with GST-GFP-NLS, sea urchin eggs were fertilized. Approximately 30 minutes after fertilization, embryos were roughly halved using a glass microinjection needle that had been pulled to be very thin. The 30-minute incubation prior to bisection allows time for perinuclear ER to accumulate. After cutting in two, the wounds generally healed quickly. The halved embryo containing the nucleus was then imaged by confocal microscopy.

###### Ectopic asymmetric division with magnetic beads

The approach to induce asymmetrically dividing embryos using magnetic beads was described previously (Salle, et al., 2019). Briefly, after a quick wash in 1 M NaCl, 1% Tween-20 solution, streptavidin magnetic beads (Nanolink-1μm; Trilink) were incubated in 2 μg/ml Atto-488-biotin PBS solution and resuspended in PBS. Beads were injected and allowed to cluster before fertilization. The bead cluster that acts as a cortical pulling cap was maintained at the cell cortex using an external magnet throughout the experiment.

##### Purified proteins

Recombinant GST-GFP-NLS was expressed and purified as previously described (Levy and Heald, 2010). The mCherry sequence was amplified from pEmCherry-C2 (a gift from Anne Schlaitz) by PCR and cloned into pMD49 (a gift from Mary Dasso, NIH) at BamHI and EcoRI, replacing the EGFP sequence to generate a bacterial expression construct for GST-mCherry-NLS (pDL94). GST-mCherry-NLS protein was purified similarly to GST-GFP-NLS (Levy and Heald, 2010). Protein stock concentrations were generally ~10 mg/ml. The Rtn4b expression construct was generated by PCR of human Rtn4b from DNASU plasmid HsCD00081743 and cloning into pET30b at EcoRI/XhoI (pDL35). GFP was cloned from pMD49 into pDL35 at EcoRV/BamHI (pDL55). Plasmid pDL55 was transformed into BL21(DE3)RIL+ *E. coli* (Stratagene) and induced at 37°C for 3 hr. Cells were resuspended in lysis buffer (50 mM Tris pH 7.6, 150 mM KCl, 5% glycerol, 10 mg/ml deoxycholic acid, 10 mM imidazole, protease inhibitors) and lysed by sonication. Lysate was rotated at 4°C for 1 hr prior to centrifugation. Cleared lysate was applied to Ni-NTA resin (Qiagen). The resin was rotated at 4°C for 1 hr, washed three times with lysis buffer, and eluted with lysis buffer containing 500 mM imidazole. Purified protein was dialyzed into 50 mM Tris pH 7.6, 150 mM KCl, 5% glycerol, 0.5 mg/ml deoxycholic acid at ~0.35 mg/ml GFP-Rtn4b. Recombinant p150CC1 was a gift from Dr. Jay Gatlin at the University of Wyoming (Gaetz and Kapoor, 2004).

##### RNA preparation

The membrane-mCherry pCS2+ was a gift from Michael Lampson (University of Pennsylvania). The H2B-RFP pCS107 plasmid (pRH199) was a gift from Rebecca Heald (University of California, Berkeley). The GFP-Rtn4b pCS107 plasmid (pDL34) was described previously (Jevtic and Levy, 2015). The calreticulin signal sequence-mCherry-KDEL cassette was PCR amplified from Addgene plasmid #55041 mCherry-ER-3 (a gift from Michael Davidson) and cloned into pCS2+ at EcoRI/XhoI (pDL107). Each plasmid was linearized with NotI except for pDL34 that was linearized with KpnI, and mRNA was expressed from the SP6 promoter using the mMessage mMachine kit (Ambion) and isolated in water. RNA stock concentrations were generally ~700-800 ng/μl.

##### Sea urchin immunostaining

Immunostaining was performed using procedures similar to those described previously (Minc, et al., 2011; Foe and von Dassow, 2008). Embryos were fixed for 70 min in 100 mM Hepes, pH 6.9, 50 mM EGTA, 10 mM MgSO_4_, 2% formaldehyde, 0.2% glutaraldehyde, 0.2% Triton X-100, and 800 mM glucose. To limit autofluorescence, samples were rinsed in PBS and placed in 0.1% NaBH_4_ in PBS made fresh 30 min before use. Samples were then rinsed in PBS and PBT (PBS plus 0.1% Triton X-100) and incubated for 24 to 48 h in rabbit anti-KDEL (Invitrogen) at 1:1000 and mouse anti-tubulin (DM1A; Sigma-Aldrich) at 1:5000 primary antibodies in PBT. After three washes of 1 h in PBT, samples were incubated for 12 h in goat anti-mouse and anti-rabbit secondary antibodies coupled with Dylight 488 and Dylight 550 (ThermoFisher Scientific) at 1:1000 in PBT. Samples were then washed three times for 1 h in PBT, incubated 15 min in PBS containing 10 μg/ml Hoechst 33342 to label DNA, transferred into 50% glycerol PBS, and finally transferred into mounting medium (90% glycerol and 0.5% N-propyl gallate PBS).

##### *Xenopus* embryo fluorescence immunocytochemistry

Immunocytochemistry of *X. laevis* blastomeres was carried out following a previously described method with a few modifications (Lee, et al., 2008). In brief, embryos were treated for 10 minutes at room temperature with 10 μg/ml proteinase K (Sigma), 2 mg/ml collagenase A (Sigma), and 20 U/ml hyaluronidase type I-S (Sigma) in 1/3× MMR to enzymatically remove the vitelline membrane. Embryos were washed three times with 1/3× MMR and any remaining vitelline membrane was removed with fine forceps. Embryos were then incubated in medium lacking calcium and magnesium (CMFM = 88 mM NaCl, 1 mM KCI, 2.4 mM NaHCO_3_, 7.5 mM Tris) to promote cell dissociation. Isolated blastomeres were fixed in freshly prepared 1x MEMFA (100 mM MOPS pH 7.4, 2 mM EGTA, 1 mM MgSO_4_, 3.7% formaldehyde) overnight at 4°C. Blastomeres were then permeabilized with 0.1% Triton-X-100 in 1× MEMFA for 2 hours at room temperature, followed by three 5-min washes in PTW buffer (0.1% Tween-20 in PBS). Blastomeres were then subjected to several 5-min washes in 100% methanol until fully dehydrated and stored at −20°C overnight. Next, surface pigments were bleached in 10% H_2_O_2_/67% methanol for 3 hours under direct light, and the blastomeres were serially rehydrated by consecutive 10 minute washes at room temperature in: 50% methanol/50% TBS; 25% methanol/75% TBS; 100% TBST (TBS + 0.1% Triton-X-100).

To reduce autofluorescence, blastomeres were incubated in an autofluorescence reducing agent (155 mM NaCl, 10 mM Tris-Cl pH 7.5, 50 mM NaBH_4_) overnight at 4°C, followed by five 10-min washes in TBST at room temperature. Blastomeres were then blocked for 2 hours at room temperature in WMBS (155 mM NaCl, 10 mM Tris-Cl pH 7.5, 10% fetal bovine serum, 5% dimethylsulfoxide). Blastomeres were incubated overnight at 4°C with primary antibodies: rabbit anti-KTN1 at 1:50 (Sigma #K1644) and mouse anti-NPC mAb414 at 1:250 (BioLegend # 902901) in WMBS. Following 5× 1 hour washes in TBST at room temperature, blastomeres were blocked overnight in WMBS at 4°C. Then the blastomeres were incubated at 4°C overnight with Alexa Fluor 488 goat anti-rabbit (Life Technologies, A-11008) and Alexa Fluor 568 goat anti-mouse (Life Technologies, A-11004) secondary antibodies at 1:250 in WMBS. Blastomeres were washed 5x 1 hour in TBST and several times in methanol (5-10 min each) until fully dehydrated. They were then cleared in 2:1 benzyl benzoate:benzyl alcohol (BBBA) for 24 hours at 4°C and imaged in custom-made imaging chambers filled with BBBA.

##### *Xenopus* egg extract, nuclear assembly, and microfluidic encapsulation

*X. laevis* metaphase-arrested crude egg extract (Good and Heald, 2018) and demembranated sperm chromatin (Hazel and Gatlin, 2018) were prepared as previously described. We fractionated crude egg extract for droplet encapsulation experiments to better visualize ER membranes and to generate a minimal membrane system lacking heavy mitochondrial membranes. Freshly prepared crude egg extract was supplemented with LPC, cytochalasin B, and energy mix and incubated on ice for 30 minutes. The extract was then subjected to a high speed clarifying spin in a Beckman TL-100 centrifuge for 22 mins at 32,000 rpm using a pre-cooled Beckman TLS-55 rotor at 4°C. This clarifying spin fractionates the crude extract into four different layers: (i) lipid layer, (ii) cytosol with light membranes (containing ER), (iii) heavy membranous layer (containing mitochondria), (iv) residual yolk platelets and insoluble materials. We collected fraction (ii) and generated an interphase-arrested extract by addition of 0.6 mM CaCl_2_ and 0.15 mg/ml cycloheximide. De novo nuclear/ER assembly was initiated by adding demembranated sperm chromatin (Chen and Levy, 2018). Fractionated egg extract was used in all droplet encapsulation experiments; crude egg extract was used in Fig. S5C-D.

Extract encapsulation experiments were performed in polydimethylsiloxane (PDMS) microfluidic devices utilizing T-junction droplet generators as previously described (Hazel, et al., 2013). PDMS (Sylgard 184, Dow Corning) microfluidic devices were replicated from a negative photoresist-on-silicon master using standard soft lithography protocols. Device depth was determined by the thickness to which photoresist was spin-coated upon the silicon wafer. PDMS replicas were trimmed and holes were punched using sharpened blunt syringe tips. Prepared devices were exposed to an oxygen plasma (Harrick Plasma) and placed in conformal contact with a glass cover slip. After baking for 10 min at 70°C, an irreversible bond was formed between the PDMS and glass, allowing sealed devices to be used as fluidic networks.

For encapsulation experiments, fractionated egg extracts were used that contained de novo assembled nuclei. To visualize nuclei, extracts were supplemented with 0.04-0.14 mg/mL recombinant GST-GFP-NLS or GST-mCherry-NLS. To visualize ER networks in fractionated extracts, CM-DiI (ThermoFisher C7000) was added at 1 μM after 15 minutes of de novo nuclear assembly. In some experiments, recombinant Rtn4b or p150CC1 was added at the indicated concentrations after 30 minutes of de novo nuclear assembly. De novo nuclear/ER assembly was carried out at 16°C for 45 minutes, with gentle flicking every 15 minutes, prior to microfluidic encapsulation in droplets. This 45-minute incubation was important because immediate encapsulation perturbed proper ER network formation in droplets. To limit liquid permeability through the PDMS walls, devices were submerged in 100 mM KCl, 0.1 mM CaCl_2_, 1.6 mM MgCl_2_, 4 mM EGTA, 50 mM sucrose, 10 mM HEPES pH 7.7 for 2 hours prior to encapsulation and during imaging. Extract and carrier oil (Pico-Surf^TM^ 2, 2% in Novec 7500, Dolomite Microfluidics, Cat No 3200282) were loaded into separate syringes and connected to their respective channel inlets via Tygon® microbore tubing (0.010” ID × 0.030” OD, Saint-Gobain Performance Plastics, Cat No AAD04091). Fluid flow to the device was established using a syringe pump (neMYSYS, Cetoni, Kent Scientific, Chemyx). Generally, oil and extract flow rates were 1-10 μl/min and 0.1-1 μl/min, respectively. Relative flow rates were adjusted to vary droplet volume. Filled devices were sealed with acrylic nail polish and droplets were incubated in microfluidic reservoirs for 3 hours at 16°C prior to confocal imaging at room temperature. This incubation period allows time for ER network formation and nuclear growth.

##### Immunofluorescence for in vitro assembled *Xenopus* nuclei and ER

For recombinant Rtn4b addition to unencapsulated extract (Fig. S5D), nuclei were assembled de novo in crude egg extract at 16°C (Chen and Levy, 2018). Recombinant 6xHis-GFP-Rtn4b was added at the indicated concentrations 30 minutes after initiating nuclear assembly. Control extract was supplemented with an equivalent volume of Rtn4b dialysis buffer. After an additional 30 minute incubation at 16°C, nuclei were fixed, spun down onto coverslips, and processed for immunofluorescence as previously described (Edens and Levy, 2014). Briefly, extract containing nuclei was mixed with 20 volumes of fix buffer (ELB, 15% glycerol, 2.6% paraformaldehyde), rotated for 15 mins at room temperature, layered over 5 mL cushion buffer (XB, 200 mM sucrose, 25% glycerol), and spun onto 12 mm circular coverslips at 1000× *g* for 15 mins at 16°C. Nuclei on coverslips were post-fixed in cold methanol for 5 mins and rehydrated in PBS-0.1% NP40. Coverslips were blocked with PBS-3% BSA overnight at 4°C, incubated at room temperature for 1 hr each with primary and secondary antibodies diluted in PBS-3% BSA, and stained with 10 μg/mL Hoechst for 5 mins. After each incubation, 6× washes were performed with PBS-0.1% NP40. Coverslips were mounted in Vectashield mounting medium (Vector Laboratories, Cat No H-1000) onto glass slides and sealed with nail polish. The primary antibody was mAb414 (BioLegend # 902901, mouse, 1:1000) that recognizes NPC FG-repeats, and the secondary antibody was a 1:1000 dilution of Alexa Fluor 568 anti-mouse IgG (Molecular Probes, A-11004). Wide-field fluorescence imaging was performed and nuclear cross-sectional areas were quantified as described previously (Edens and Levy, 2014). To validate the proper incorporation of recombinant Rtn4b into ER membranes (Fig. S5C), similar de novo nuclear assembly was performed, and 67 nM 6xHis-GFP-Rtn4b was added 30 minutes after initiating nuclear assembly. Then, 1 μM CM-DiI was added to visualize the ER and a 1:1000 dilution of Alexa Fluor 488 conjugated 6x-His Tag Monoclonal Antibody (Invitrogen, MA1-21315-A488) was added to visualize His-tagged Rtn4b. After an additional 30 minute 16°C incubation, a small aliquot of the assembly reaction was squashed between a glass slide and coverslip and sealed with VALAP. ER networks were allowed to form for 45 minutes and then imaged by wide-field fluorescence microscopy.

##### *Xenopus* microscopy

For Fig. S5C-D, wide-field microscopy was performed with a fluorescence microscope (BX51; Olympus) using the following objectives: UPLFLN 20x (NA 0.50, air; Olympus), UPLFLN 40x (NA 0.75, air; Olympus), and UPLANAPO 60× (NA 1.20, water; Olympus). Images were acquired with a QIClick Digital charge-coupled device (CCD) camera, mono, 12-bit (model QIClick-F-M-12) using cellSens software (Olympus). Images for measuring fluorescence staining intensity were acquired using the same exposure times. Total fluorescence intensity and cross-sectional nuclear area were measured from the original thresholded images using MetaMorph software (Molecular Devices).

All other *Xenopus* imaging was performed using a spinning-disk confocal microscope based on an Olympus IX81 microscope stand equipped with a five line LMM5 laser launch (Spectral Applied Research) and Yokogawa CSU-X1 spinning-disk head. Confocal images were acquired with an EM-CCD camera (ImagEM, Hamamatsu). Z-axis focus was controlled using a piezo Pi-Foc (Physik Instrumentes), and multiposition imaging was achieved using a motorized Ludl stage. Olympus objectives included: PLanApo 60× (NA 1.35, oil), UPLanSApo 40x (NA 1.25, silicon oil), UPLFLN 40× (NA 0.75, air). Metamorph software was used to control system components and for image acquisition. Images for measuring fluorescence intensity were acquired using the same exposure times.

To image fixed dissociated *Xenopus* blastomeres (Fig. 6A-B), multiple z-sections were acquired through each blastomere for both KTN1 and mAb414 staining (0.4 μm z-distance for stage 8 and 0.3 μm z-distance for stage 9 and 10). Confocal z-stacks were 3D reconstructed using Metamorph. Since KTN1 exclusively stains perinuclear ER sheets, we thresholded the perinuclear ER and calculated the volume based on isosurface rendering. Nuclear surface area was extrapolated from CS area measurements based on mAb414 staining and assuming a sphere, as validated (Fig. S1B) and previously described (Vukovic, et al., 2016b; Jevtic and Levy, 2015; Edens and Levy, 2014; Levy and Heald, 2010).

To image extract droplets (Fig. 6C-F and S5A-B), live nuclei in droplets were visualized with GFP-NLS and ER was visualized with DiI. Z-sections 1.5 μm thick were acquired through each droplet in both fluorescence channels at room temperature.

#### QUANTIFICATION AND STATISTICAL ANALYSIS

##### Image analysis

Images were processed and analyzed with Fiji (ImageJ; National Institutes of Health) and assembled in Photoshop and Illustrator (Adobe). For publication, images were cropped and pseudocolored using Fiji, but were otherwise unaltered.

To quantify nuclear size, the maximum cross-sectional (CS) nuclear area was measured using GFP-NLS or mCherry-NLS and nuclear surface area was extrapolated by multiplying by 4. We validated this method for estimating nuclear surface area by showing that actual nuclear volume measured from confocal z-stacks matches well with nuclear volume estimated from the CS area assuming a sphere (Fig. S1B), showing that the nuclear shape approximates a sphere. This is consistent with previous data from various systems showing that CS nuclear area accurately predicts total nuclear surface area and volume as measured from confocal z-stacks (Vukovic, et al., 2016b; Jevtic and Levy, 2015; Edens and Levy, 2014; Levy and Heald, 2010). We note in the figure legends whether nuclear size was quantified from confocal or wide-field imaging. For confocal imaging large numbers of z-planes were acquired, whereas only a few z-planes were acquired for wide-field imaging. As a consequence, we noticed slight deviations in absolute nuclear size values between these two different acquisition methods.

Cell volumes were extrapolated from cell CS areas quantified from membrane-mCherry localized at the plasma membrane, approximating cells as spheres (Fig. S1C). To measure nuclear import, average nuclear GFP-NLS intensity was measured and corrected for nuclear volume to obtain the total intranuclear GFP-NLS signal. These intensity values were plotted as a function of time and the initial slope was used to calculate the initial nuclear import rate. Import rates were normalized to the cytoplasmic GFP-NLS signal in the preceding mitosis.

Perinuclear ER volume was calculated based on perinuclear KDEL or DiI signal (see Fig. S4B for details). To quantify perinuclear ER accumulation (Fig. 4A and 4C), we measured the mean KDEL intensity in a 15-μm wide ring around the nucleus. First, a circle was drawn around the nucleus and the total KDEL intensity within that circle was quantified. A second circle with radius 15 μm greater than that of the first circle was drawn around the perinuclear region and the total KDEL intensity within that circle was quantified. The total KDEL intensity in the perinuclear ring was calculated by subtracting the total KDEL intensity value of the first circle from the second. The perinuclear ER accumulation index was calculated by dividing the mean KDEL intensity within this 15-μm wide perinuclear ring by the mean KDEL intensity of the whole cell. To quantify the radial distribution pattern of the ER (Fig. 4C), we plotted the mean KDEL intensity in 2.5 μm wide concentric rings, starting from the nuclear envelope and radiating towards the cell periphery. The area and total KDEL intensity were quantified for the first 2.5 μm wide perinuclear ring closest to the NE as described above for Fig. 4A and 4C. This process was repeated for up to fourteen 2.5 μm-wide concentric rings around the first perinuclear ring, until their radial distribution reached the cell boundary. Mean KDEL intensities were calculated by dividing the total intensity of each ring by its area.

In vitro droplet volumes (Fig. 6C-F) were calculated from droplet diameter measurements. To measure nuclear size, we first compared nuclear volumes quantified from 3D reconstructed confocal z-stacks to nuclear volumes extrapolated from maximum nuclear CS areas assuming spherical nuclei, finding these values agreed within 3% on average (data not shown). This is consistent with previous data showing that CS nuclear area accurately predicts total nuclear surface area and volume as measured from confocal z-stacks (Vukovic, et al., 2016b; Jevtic and Levy, 2015; Edens and Levy, 2014; Levy and Heald, 2010) (Fig. S1B). Based on this validation, we measured CS nuclear areas from original thresholded images using Metamorph and then extrapolated surface area by multiplying by 4. Perinuclear ER volume was calculated based on perinuclear DiI signal as described in Figure S4B.

##### Modeling of nuclear growth kinetics

The equation for nuclear stage *N*_*i*_(t) can be solved analytically, leading to:

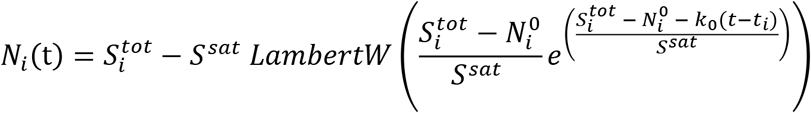

In which 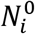 is the initial nucleus size at stage *i* (obtained experimentally) and *t*_*i*_ is the initial time. To fit the four parameters, we used Matlab’s *fminsearch* function to minimize the relative error or nuclear size:

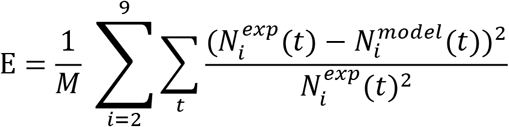

In which *M* = 113 is the total number of experimental points. We found the parameters *k*_0_ = 52.07 *min*^−1^, λ = 0.171, *α* = 0.546, *S*^*sat*^ = 241.9 μm^2^. This results in a fitting error of 0.36 % (defined such that 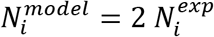 would yield an error of 100%). The fitting scripts, together with the necessary experimental data, are available online: https://github.com/SergeDmi/NuclearGrowth.

We note that in this model we did not fit the data from the one-cell stage because the progressive accumulation of the ER on to the nuclear surface introduces a time-dependent input for the ER amount, which differs from other stages (Fig. 4A).

##### Statistical analysis

Averaging and statistical analysis were performed for independently repeated experiments. Where indicated, nuclear size and intensity measurements were normalized to controls. Two-tailed Student’s t-tests assuming equal variances and confidence ellipses were calculated using Excel to evaluate statistical significance. The p-values, number of embryos examined, number of nuclei and cells quantified, and error bars are denoted in the figure legends.

## KEY RESOURCES TABLE

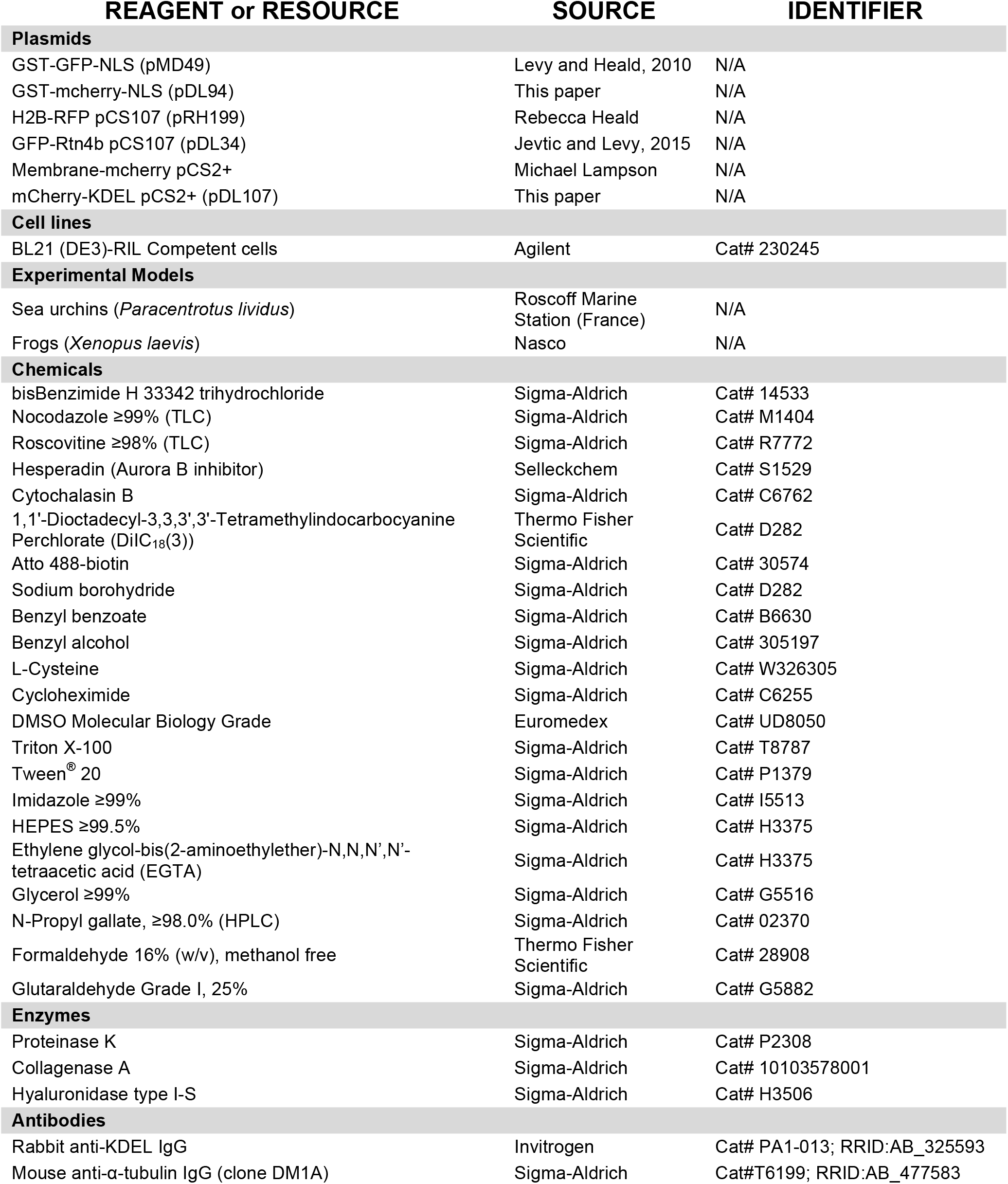
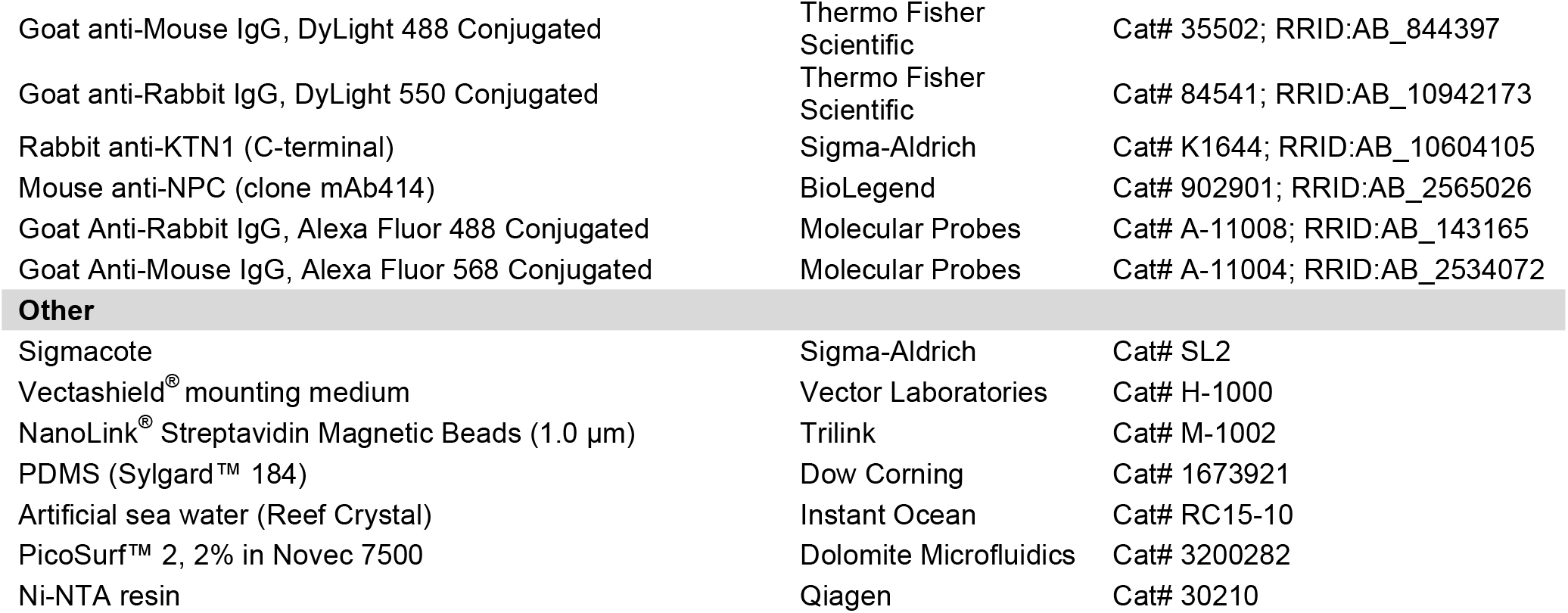

