## Supplemental Information for "The perinuclear ER scales nuclear size independently of cell size in early embryos"

### SUPPLEMENTAL FIGURE LEGENDS

#### Figure S1

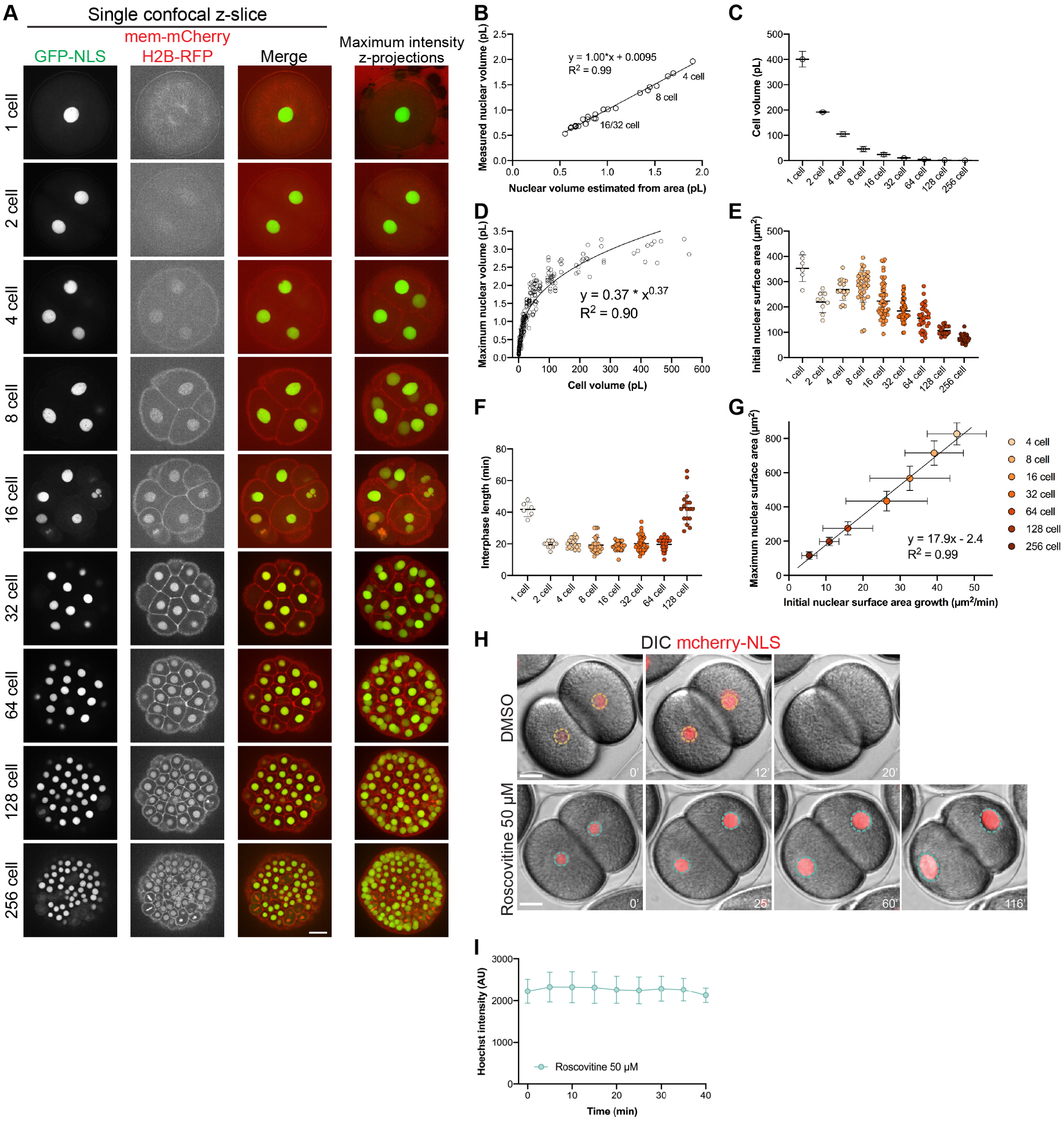


**Figure S1. Additional data related to nuclear size scaling in sea urchin embryos. (A-G)** Additional quantification related to Fig. 1A-E. **(A)** Representative images from different embryos and time-lapses. Brightness and contrast were adjusted to better visualize cell boundaries and nuclei. **(B)** Nuclear volumes calculated from confocal z-stacks were compared to nuclear volumes estimated from maximum cross-sectional

(CS) nuclear areas assuming roughly spherical nuclei (n=24). The data indicate that CS nuclear area is a good proxy for nuclear volume and nuclear surface area. This is consistent with previous data from a variety of different systems showing that CS nuclear area accurately predicts nuclear surface area and volume as measured from confocal z-stacks (Vukovic, et al., 2016; Jevtic and Levy, 2015; Edens and Levy, 2014; Levy and Heald, 2010). **(C)** Maximum cross-sectional cell areas were measured based on membrane-mCherry localized at the plasma membrane and used to extrapolate cell volume assuming roughly spherical cells. On average, these measured values matched well with calculations assuming blastomere volumes are halved with each cell division. **(D)** Maximum nuclear volume is plotted as a function of cell volume. These are the same data from Fig. 1D plotted here on linear x- and y-axes and fit to a power series. **(E)** Initial nuclear surface areas were measured from the time when intranuclear GFP-NLS signal was first visible after nuclear assembly. Apparent reductions in initial nuclear size after the 16-cell stage may be due to the reduction in nuclear growth rates. After nuclear assembly occurs, several minutes pass before the acquisition of the first post-mitotic image for a given cell. Since nuclear growth is faster in the early embryo, more nuclear growth will happen during this window, potentially explaining why the initial nuclear size appears larger earlier in development. The presence of karyomeres during early nuclear formation (Movies 1-3) also complicates initial nuclear size measurements. **(F)** Interphase length was determined from time-lapses as the time between the first visible intranuclear GFP-NLS signal to the time of NE breakdown when intranuclear GFP-NLS signal dissipated. **(G)** Average maximum nuclear surface area is plotted as a function of initial nuclear growth rate for different developmental stages and fit to a linear regression. **(H)** Embryos were microinjected with GST-mCherry-NLS protein and treated with 50 μM roscovitine or an equal volume of DMSO at the 2-cell stage. Nuclear surface areas were extrapolated from CS areas at one-minute intervals based on wide-field imaging. Representative images are shown, and the quantification is in Fig. 1G. **(I)** One-cell embryos were treated with 50 μM roscovitine and DNA was stained with Hoechst. Hoechst staining intensity was quantified at 5-minute intervals (n=15).

Error bars represent SD. Scale bars: 20 μm.

**Figure S2**

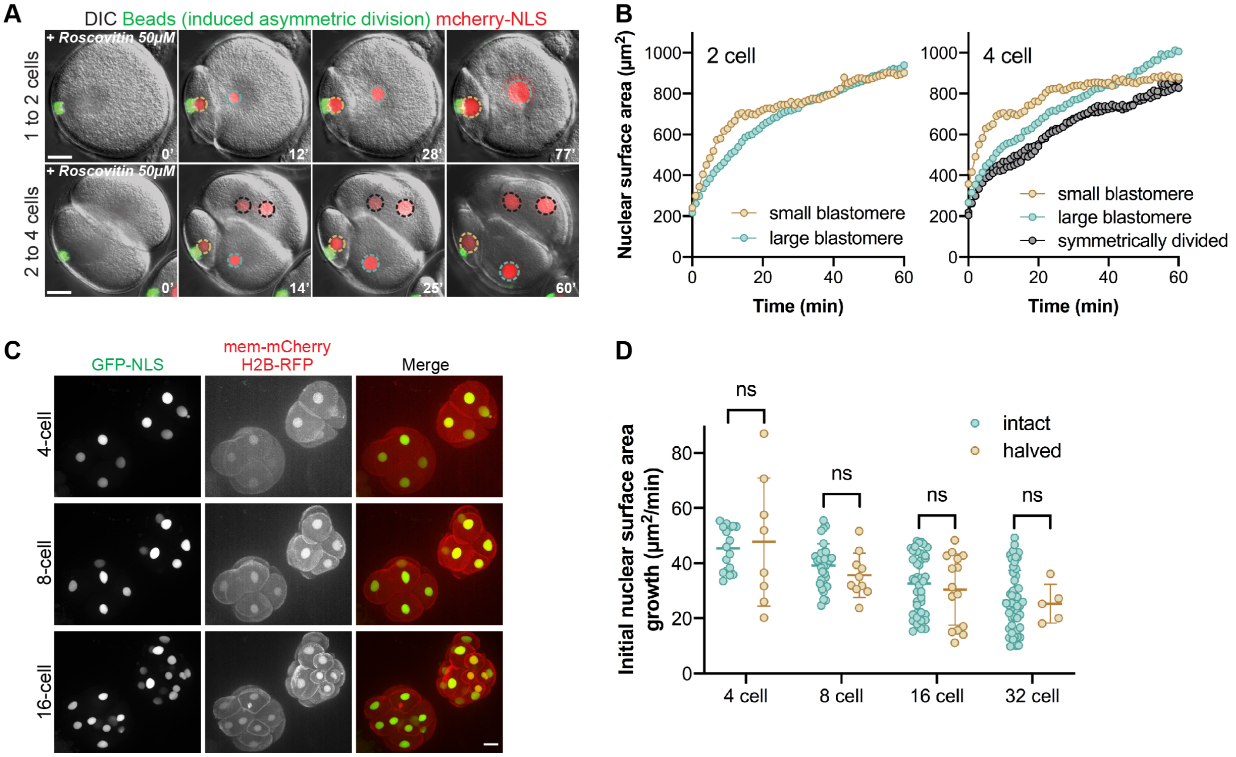


**Figure S2. Additional data related to nuclear size scaling in asymmetrically divided and halved embryos. (A-B)** Asymmetric cell divisions were induced at the first or second cleavage as described in Fig. 2E-F in the presence of 50 μM roscovitine. Nuclear surface areas extrapolated from CS areas were quantified at 1-minute intervals based on wide-field imaging. Wide-field imaging was performed with a limited number of z-planes so these size measurements should not be compared to data obtained from confocal imaging. **(C-D)** Additional quantification related to Fig. 3A-C. **(C)** Representative images of halved embryos are shown at different developmental stages as maximum intensity z-projections. Brightness and contrast were adjusted to better visualize cell boundaries and nuclei. **(D)** Initial nuclear growth rates were calculated based on the first 3-5 time points of each nuclear growth curve. Intact embryo data are the same shown in Fig. 1E.

Error bars represent SD. ns, not significant. Scale bars: 20 μm.

### Figure S3

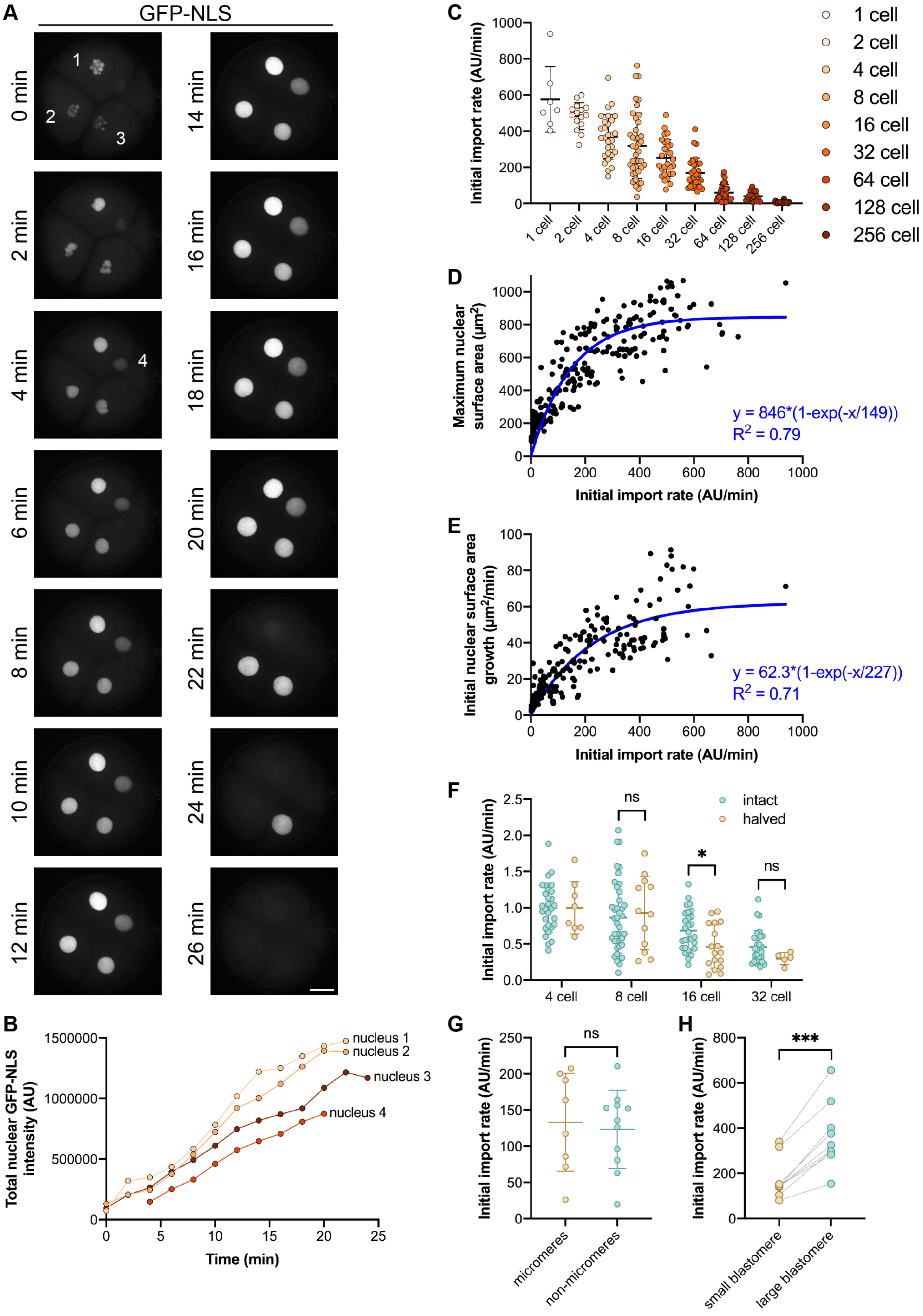
A
GFP-NLS
0 min
2 min
4 min
6 min
8 min
10 min
12 min
14 min
16 min
18 min
20 min
22 min
24 min
26 min
1
2
3
4
B
Total nuclear GFP-NLS intensity (AU)
1500000
1000000
500000
0
0
5
10
15
20
25
Time (min)
nucleus 1
nucleus 2
nucleus 3
nucleus 4
C
Initial import rate (AU/min)
1000
800
600
400
200
0
1 cell
2 cell
4 cell
8 cell
16 cell
32 cell
64 cell
128 cell
256 cell
D
Maximum nuclear surface area (μm²)
1000
800
600
400
200
0
0
200
400
600
800
1000
Initial import rate (AU/min)
y = 846*(1-exp(-x/149))
R² = 0.79
E
Initial nuclear surface area growth (μm²/min)
100
80
60
40
20
0
0
200
400
600
800
1000
Initial import rate (AU/min)
y = 62.3*(1-exp(-x/227))
R² = 0.71
F
Initial import rate (AU/min)
2.5
2.0
1.5
1.0
0.5
0.0
intact
halved
ns
*
ns
4 cell
8 cell
16 cell
32 cell
G
Initial import rate (AU/min)
250
200
150
100
50
0
ns
micromeres
non-micromeres
H
Initial import rate (AU/min)
800
600
400
200
0
***
small blastomere
large blastomere

**Figure S3. Nuclear import kinetics in sea urchin embryos. (A-E)** Experiments described in Figure 1A-E were analyzed for nuclear import. **(A)** Maximum intensity z-projections of the same embryo over time. **(B)** Intranuclear GFP-NLS intensity was quantified for the nuclei shown in (A) (see Methods for details). **(C)** Initial nuclear import rates were calculated based on the first 3-7 time points of each nuclear import curve and normalized to the cytoplasmic GFP-NLS signal in the preceding mitosis (see Methods for details). Initial nuclear import rates are plotted for different developmental stages. Nucleus number: n=7 (1-cell), n=15 (2-cell), n=29 (4-cell), n=41 (8-cell), n=28 (16-cell), n=31 (32-cell), n=32 (64-cell), n=18 (128-cell), n=29 (256-cell). **(D)** For different developmental stages, maximum nuclear surface area is plotted as a function of the initial nuclear import rate. The blue curve shows the one-phase association fit. Nucleus number: n=7 (1-cell), n=15 (2-cell), n=29 (4-cell), n=41 (8-cell), n=28 (16-cell), n=33 (32-cell), n=32 (64-cell), n=18 (128-cell), n=29 (256-cell). **(E)** For different developmental stages, the initial nuclear growth rate is plotted as a function of the initial nuclear import rate. The blue curve shows the one-phase association fit. Nucleus number: n=7 (1-cell), n=13 (2-cell), n=17 (4-cell), n=17 (8-cell), n=26 (16-cell), n=33 (32-cell), n=32 (64-cell), n=16 (128-cell), n=29 (256-cell). **(F)** Initial nuclear import rates of GFP-NLS were measured for halved embryos described in Fig. 3A-C and normalized to the 4-cell stage data. n=8 (4-cell), n=12 (8-cell), n=17 (16-cell), n=5 (32-cell). Intact embryo data are the same shown in (C). **(G)** Initial nuclear import rates of GFP-NLS were measured for micromeres versus non-micromeres at the 16-cell stage (also see Fig. 2A-D). n=8 (micromeres), n=11 (non-micromeres). **(H)** Initial nuclear import rates of GFP-NLS were measured for small and large blastomeres generated by induced asymmetric cell divisions (also see Fig. 2E-F). n=9 (small blastomeres), n=9 (large blastomeres).

Error bars represent SD. ***, $p<0.005$; *, $p<0.05$; ns, not significant. Scale bars: 20 μm.

**Figure S4**

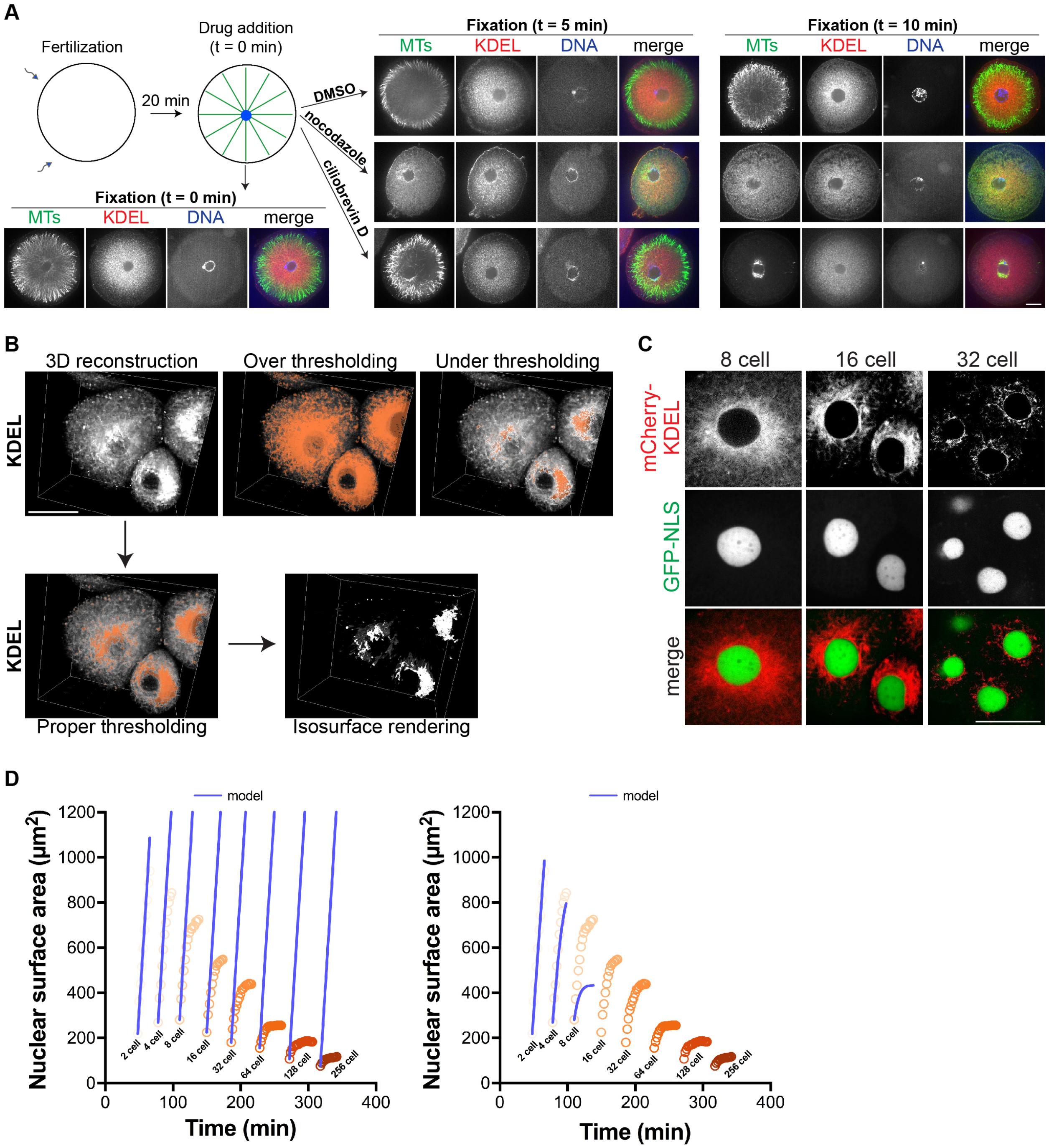


**Figure S4. Additional data related to ER measurements. (A)** Twenty minutes after fertilization, embryos were fixed or treated with 20 μM nocodazole, 50 μM ciliobrevin D, or DMSO as a control. Embryos were fixed 5 or 10 minutes later and immunostained with anti-KDEL and anti-tubulin antibodies. Relevant quantifications are shown in Fig. 4C. Note that the KDEL images shown here 10 minutes after drug addition are the

same shown in Fig. 4C. **(B)** Method to quantify perinuclear ER volume in sea urchin embryos. Live imaging of ER was performed in embryos microinjected with mCherry-KDEL mRNA or stained with DiI. Confocal z-slices were acquired through the embryo with a 3-μm step size. The z-stacks were 3D reconstructed using Metamorph software. Perinuclear ER was defined by inclusive thresholding for each cell in 3D. Because KDEL and DiI are general ER markers, we performed manual thresholding to include the strong perinuclear ER signal and exclude the dimmer cortical ER signal. The first panel shows a representative 3D reconstructed image of a portion of a 16-cell stage embryo. To the right are examples of over thresholding and under thresholding. The second row of panels shows proper thresholding followed by isosurface rendering of the properly thresholded image. 3D voxel volume of the perinuclear ER was quantified based on this isosurface. **(C)** Sea urchin eggs were microinjected with GST-GFP-NLS protein and mRNA encoding mCherry-KDEL prior to fertilization. Representative confocal images at different developmental stages are shown. Contrast was similarly adjusted in each image to highlight the brighter perinuclear ER signal. See Fig. 5B-C for the relevant quantification. **(D)** (Left) Results of the model using no input from the ER ($\alpha=0$), overlaid on experimental data. (Right) Results of the model using a linear relation between ER volume and surface available for nuclear growth from the ER ($\alpha=1$), overlaid on experimental data. Note that after the 8-cell stage, no material is left for the nuclei to reform or grow, so all model curves are at 0.
Scale bars: 20 μm.

Figure S5

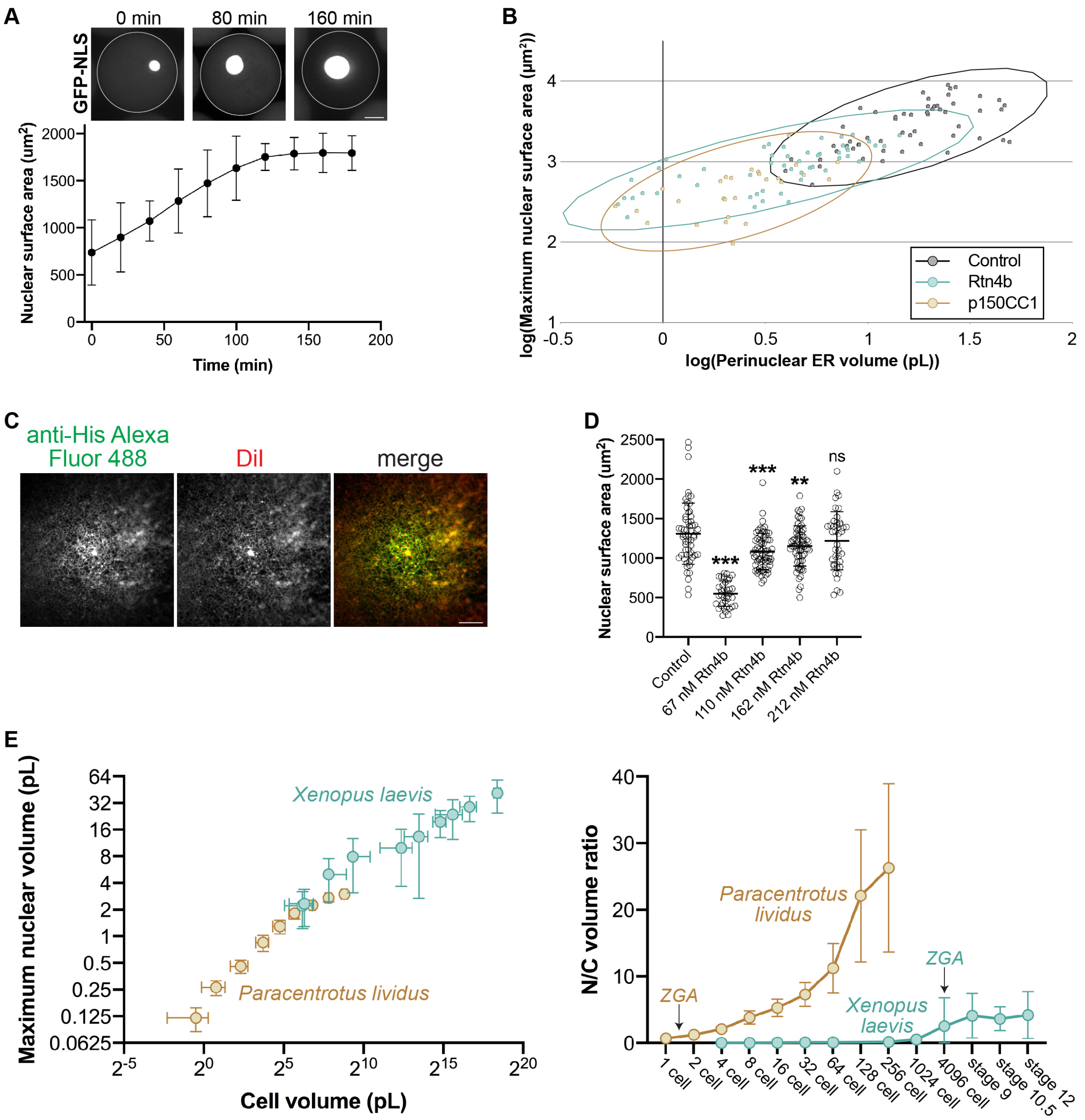


**Figure S5. Additional data related to *Xenopus in vitro* experiments and comparisons of nuclear size scaling in sea urchins and *Xenopus*. (A)** Nucleus formation was induced in fractionated interphase *X. laevis* egg extract supplemented with GST-GFP-NLS protein. Extract and nuclei were encapsulated in 80-90 µm diameter droplets using microfluidics. Once intranuclear GFP-NLS was apparent indicating formation of an intact NE, confocal images were acquired at 20-minute intervals. At each time point nuclear CS area was measured for 8-12 nuclei (10 nuclei on average), and nuclear surface area was extrapolated for the nearly spherical nuclei. Nuclear growth is plotted as a function of time. **(B)** Nuclear surface area as a function of perinuclear ER volume is plotted for the droplet data shown in Fig. 6F with 95% confidence ellipses. **(C)** To validate that recombinant Rtn4b added to *Xenopus* extract properly incorporates into reconstituted ER, crude interphase egg extract was supplemented with 1 μM DiI and 67 nM recombinant Rtn4b. To label the His-tagged Rtn4b, an Alexa Fluor 488 labeled anti-His antibody was added (see Methods for details). Representative images are shown. The reticular localization of Rtn4b co-localizing with DiI demonstrates its proper integration into the ER. **(D)** To determine the concentration of exogenously added Rtn4b that most limits nuclear growth, we assembled nuclei de novo in crude *X. laevis* egg extract and titrated in recombinant Rtn4b at the indicated concentrations. After a 30-minute incubation, nuclear CS areas were quantified based on NPC staining and extrapolated to surface areas (see Methods for details). 37-66 nuclei were quantified per condition (53 nuclei on average). Because nuclei were smallest with 67 nM added Rtn4b, we used this concentration of Rtn4b throughout the study. **(E)** Nuclear volume is plotted as a function of cell volume for different urchin and *Xenopus* developmental stages. Urchin data are from Fig. 1D. Frog data were published previously (Jevtic and Levy, 2015). N/C volume ratios were calculated for individual urchin blastomeres and averaged for a given stage based on the data in Fig. 1D. *X. laevis* N/C volume ratios were published previously (Jevtic and Levy, 2015). The timing of zygotic genome activation (ZGA) in each species is noted based on published reports (Poccia, et al., 1985; Newport and Kirschner, 1982; Nemer, 1963).

### MOVIE LEGENDS

**Movie 1. Urchin nuclear size scaling from 1- to 32-cell stage.** Sea urchin eggs were microinjected with GST-GFP-NLS protein (green) prior to fertilization. Confocal imaging was performed at 1-minute intervals for 200 minutes, and maximum intensity projections are shown. In early cleavages, assembly of import-competent nuclei occurs around individual or small clusters of chromosomes generating karyomeres that ultimately fuse to form a single nucleus. Karyomeres have been observed in a variety of species (Samwer, et al., 2017; Abrams, et al., 2012; Lemaitre, et al., 1998).

**Movie 2. Urchin nuclear size scaling from 2- to 16-cell stage.** Sea urchin eggs were microinjected with GST-GFP-NLS protein (green) and mRNA encoding membrane-mCherry (red) and H2B-RFP (red) prior to fertilization. Confocal imaging was performed at 2-minute intervals for 118 minutes, and maximum intensity projections are shown. In early cleavages, assembly of import-competent nuclei occurs around individual or small clusters of chromosomes generating karyomeres that ultimately fuse to form a single nucleus. Karyomeres have been observed in a variety of species (Samwer, et al., 2017; Abrams, et al., 2012; Lemaitre, et al., 1998).

**Movie 3. Urchin nuclear size scaling from 32- to 256-cell stage.** Sea urchin eggs were microinjected with GST-GFP-NLS protein (green) and mRNA encoding membrane-mCherry (red) and H2B-RFP (red) prior to fertilization. Confocal imaging was performed at 2-minute intervals for 134 minutes, and maximum intensity projections are shown.

**Movie 4. Urchin nuclear growth in micromeres versus macromeres.** Sea urchin eggs were microinjected with GST-GFP-NLS protein (green) and mRNA encoding membrane-mCherry (red) and H2B-RFP (red) prior to fertilization. Confocal imaging was performed at 2-minute intervals for 24 minutes, and maximum intensity projections are shown. One micromere and one macromere were cropped at the 16-cell stage.

**Movie 5. Urchin nuclear size scaling in asymmetrically dividing embryo.** Sea urchin embryos were microinjected with GST-mCherry-NLS protein (red) and magnetic beads (green). An external magnet was used to induce asymmetric divisions. Wide-field DIC and fluorescence imaging was performed at 1 minute intervals for 186 minutes.

**Movie 6. Urchin nuclear size scaling in halved embryos.** Sea urchin eggs were microinjected with GST-GFP-NLS protein (green) and mRNA encoding membrane-mCherry (red) and H2B-RFP (red), fertilized, and bisected with a glass pipet ~30 min post-fertilization. Confocal imaging was performed at 5-minute intervals for 90 minutes, and maximum intensity projections are shown.

**Movie 7. Perinuclear ER accumulation and partitioning during urchin development.** Sea urchin eggs were stained with DiI and fertilized. Confocal imaging was performed at 3-minute intervals for 174 minutes.
